# Migration and standing variation in vaginal and rectal yeast populations in recurrent vulvovaginal candidiasis

**DOI:** 10.1101/2023.07.19.549743

**Authors:** Abdul-Rahman Adamu Bukari, Rebekah J. Kukurudz-Gorowski, Alexia de Graaf, Devin Habon, Beamlak Manyaz, Yana Syvolos, Aruni Sumanarathne, Vanessa Poliquin, Aleeza Gerstein

## Abstract

Vulvovaginal candidiasis is one of the most common fungal infections. Most are successfully treated with antifungal drugs, yet ∼8% lead to recurrent vulvovaginal candidiasis ("RVVC"). Previous research found closely-related isolates within vaginal and rectal populations. However, their methods preclude assessing fine-scale relationships among closely related isolates and measuring genetic variation, a fundamental property with evolutionary potential implications. To address this gap, we isolated 12 vaginal and 12 rectal yeast isolates during symptomatic relapse from four individuals with a history of RVVC. Three had *Candida albicans* infections, while the fourth had *Nakaseomyces glabratus*. All isolates were whole-genome sequenced and phenotyped. The isolates were placed into global phylogenies built from short-read WGS data, including an updated *N. glabratus* tree with over 500 isolates. Genotypic and phenotypic analyses were consistent with migration between sites. There was little phenotypic diversity for drug response and no consistent difference between isolates from different sites for invasive growth. Although there are few comparables, *C. albicans* nucleotide diversity was similar to most commensal oral and rectal populations, while *N. glabratus* was similar to some bloodstream infections (though higher than others). Single nucleotide changes drove diversity; no aneuploidies were found, and only a single loss-of-heterozygosity tract on chr1L varied among isolates from one participant. This study provides baseline measurements and describes techniques to quantify within-population diversity in fungal microbes. We highlight a need for comparable studies that use the same sampling effort and analysis methods to understand the interplay in shaping fungal microbial communities in important contexts.

**Author Summary:** Recurrent vaginal yeast infections are relatively common, and we do not understand why some people experience these chronic infections when many others have a single infection that is successfully treated and cleared. Many open questions remain about the basic biology of the yeast populations involved. We quantified diversity using modern sequencing technology within vaginal and rectal yeast populations from four individuals with a history of recurrent yeast infections experiencing symptoms. Three participants had a *Candida albicans* infection (the most common causative species), while the fourth had a *Nakaseomyces glabratus* infection (the second most common and increasingly implicated). We found that vaginal and rectal isolates were closely related, indicating the same population is present at the two sites. Surprisingly, we found that diversity was similar to the yeast populations found at other body sites in healthy people. Our study highlights a critical need for additional studies following the same methods in different contexts to better understand the fungal microbial populations in our bodies.

## Introduction

Vulvovaginal candidiasis (VVC, colloquially "yeast infection") is a lower female reproductive tract mucosal infection. It is very common, affecting approximately 75% of people defined female at birth at least once in their lives (1–3). The disease burden of VVC results in global annual treatment costs of ∼ 1.8 billion USD (4, 5), with a loss in productivity in high-income countries of ∼ 14 billion USD (4). Treatment involves topical or oral antifungal medication, which is effective at symptom abatement in most cases. However, ∼ 8% of individuals with VVC experience recurrence (RVVC), defined as three or more symptomatic episodes a year (4, 5). Approximately half of all people with RVVC have no identifiable risk factors (6), signifying the need for additional studies on the biological basis of this chronic condition.

*Candida albicans* is responsible for 50-90% of VVC and RVVC cases (collectively, R/VVC) (7–11). *Nakaseomyces glabratus* (formerly *Candida glabrata*) is the second most prevalent cause, globally attributed to ∼ 8% of cases (12–14). Here, we collectively refer to these species using the colloquial term "yeast," which reflects a shared morphology while acknowledging our current understanding of their phylogenetic relationships and the recent official renaming of *N. glabratus* (15, 16). To be consistent with clinical practice, we continue to use the R/VVC abbreviations while noting that "candidiasis" does not reflect the updated genera names. Treatment recommendation for R/VVC differs by species, as many isolates from *N. glabratus* (and other non-*albicans* pathogenic yeast species) have intrinsic resistance to the azole antifungal fluconazole that is commonly used to treat RVVC (17).

Understanding the etiology of R/VVC is complicated in part since yeast can exist as a common commensal member of the vaginal microbiota without causing symptomatic VVC. Identification of vaginal yeast in the absence of symptoms does not warrant treatment, and self-diagnosis due to symptoms has also been shown to be problematic, as other gynecological infections and dermatologic conditions have overlapping symptoms (18). Approximately 70% of healthy women were found to have vaginal yeast at a single time point through an amplicon-based study (19). Genetic typing and phylogenetic strain analyses in *C. albicans* have repeatedly found no strict phylogenetic differentiation between commensal and pathogenic strains. However, some phylogenetic clades are potentially over-represented by commensal strains and those causing superficial disease (20, 21).

Relapse in RVVC, i.e., return of symptoms, could theoretically be due to either incomplete eradication of the vaginal yeast population or complete vaginal eradication followed by re-colonization (22). There are decades of studies that have sought to understand the etiology of RVVC, as the answer has potential implications for improving treatment and reducing or eliminating symptom recurrence. The GI tract has been suggested as a possible endogenous source population. A study in 1979 that treated RVVC patients with oral nystatin to reduce the resident GI population found that it did not decrease the time to recurrence (23). Furthermore, studies examining yeast colonization of the GI tract through feces or rectal swabs during symptomatic recurrence find that not all participants are culture-positive (23–29). However, this does not necessarily preclude that a small GI population is present in all individuals (below the culture detection limit in some), which could act as an endogenous reintroduction source under the right host conditions. Examining the diversity of strains at different body sites and among recurrent infections can potentially differentiate between relapse scenarios. If vaginal isolates are closely related to GI tract/rectal isolates but less diverse, that would be consistent with reintroduction. If vaginal isolates sampled at different time points consistently have the same genotype that is not present in the GI tract/rectal isolates, this would be consistent with incomplete eradication. Looking at the level of diversity and relationships among isolates acquired within a single time point ("standing variation") at different body sites can also help us understand the adaptive potential and migration dynamics of yeast populations within the body.

Recent studies that seek to map phylogenetic relationships among isolates typically use multilocus sequence typing (MLST) (11, 30, 31). The overarching results are consistent with a role for the maintenance of genotypes (i.e., the same or very similar genotype is recovered from different body sites and within the vagina between successive symptomatic episodes), alongside an occasional opportunity for novel genotypes to arise (i.e., through reinfection, sometimes referred to as strain turnover). However, previous studies do not necessarily rule out a role for standing genetic variation to have been consistently present yet undetected, i.e., relapse masquerading as reinfection, as typically only one or a small number of isolates were examined at a given time. Only a single study sequenced two vaginal isolates from different time points through short-read whole genome sequencing (WGS) (32) , and no previous studies have used WGS to compare multiple isolates from the same time point. WGS is required to accurately assess relatedness, diversity, and the potential source of genetic novelty. WGS removes the need to rely on a single or small number of markers, which could over or underinflate the actual level of diversity. For example, Sitterlé et al. showed that while MLST revealed occasional differences among *C. albicans* oral isolates collected from three healthy individuals, whole-genome sequencing revealed that the three examined isolates were closely related in each case (33).

Standing genetic variation of yeast populations has only been quantified in a handful of contexts. Two studies in *C. albicans* that whole genome sequenced 3-6 oral and rectal isolates from six healthy individuals found that although isolates were closely related in most cases, they differed by numerous single nucleotide polymorphisms, primarily resulting from short-range loss-of-heterozygosity tracts (21, 33). Two individuals were, however, simultaneously colonized with oral isolates from different clades (21). A single *N. glabratus* study sequenced up to ten isolates from nine patients with bloodstream candidemia (34); a pairwise SNP analysis was also consistent with closely related isolates. Variation in RVVC has yet to be quantified through WGS; hence, whether it is similar to commensal populations is unknown. It also needs to be determined how many isolates must be sequenced to capture population diversity accurately.

Past studies in other contexts have shown that population genetic analysis can be conducted with data from four to six individuals when including many variants (as found in data from short-read WGS data, (35–37). These studies were conducted in sexually reproducing organisms; however, how many yeast isolates are required to accurately access the population level of diversity remains unexplored.

Here we build on previous work by using short-read WGS paired with high-throughput phenotyping to quantify vaginal and rectal standing variation in participants with a history of RVVC when high vaginal and rectal yeast population sizes were observed. We compared our results to the few comparable studies that conducted short-read WGS of contemporaneous yeast isolates from other contexts and statistically evaluated the minimum number of isolates required to measure genetic variation accurately. We obtained isolates from the same time point from four individuals with a history of RVVC in Winnipeg, Canada. Three participants had *C. albicans* infections, and one had *N. glabratus*. We collectively refer to these isolates as the "THRIVE-yeast" isolates, following the name of our local umbrella research program that studies The Host-microbial Relationships and Immune function in different Vaginal Environments ("THRIVE," http://www.mthrive.ca). From all individuals, we found a complete phylogenetic overlap of vaginal and rectal isolates and no consistent difference in phenotypes, consistent with high levels of migration. We found no evidence that diversity in populations from the two sites was different; levels of standing genetic variation were generally similar to what has been observed among commensal oral and rectal populations in *C. albicans* and some bloodstream infections in *N. glabratus*. This suggests that despite frequent population bottlenecks caused by drug treatment, vaginal yeast diversity is maintained or rapidly restored.

## Materials and Methods

### Clinical isolates

Clinical samples were collected from a single time point from four non-pregnant female participants aged 18-50 years. Seventeen consenting female participants who attended a clinic for individuals with a history of recurrent vulvovaginal candidiasis (RVVC) in Winnipeg, Canada, were sampled at the clinic for possible inclusion in this study. Vaginal and rectal swabs were acquired from all participants. Swabs were kept at -4 ℃ and on ice during transport and processed within 5 h of acquisition. Swabs were then agitated for ∼30 s in 1 mL PBS, and a dilution series (1, 1:10, 1:100) was conducted. 100 µL from each of the three dilutions was spread onto SDA and chromogenic *Candida* agar plates. Plates were incubated for 48 h at 30 `°`C. Of the seventeen participants, vaginal colonies were acquired from ten. Of those, six were also positive for rectal isolates. Our goal in this study was to compare standing genetic and phenotypic variation among vaginal and rectal populations, as well as to statistically determine the number of isolates required to infer within-population diversity at a single time point accurately. To meet these goals, we focused our efforts on the four participants who had high vaginal and rectal populations (> 1 × 10^3^ CFU/mL of swab elute). Twelve vaginal and 12 rectal isolates were haphazardly isolated from each participant from dilution plates that had clear margins between colonies. Colonies were suspended in 1 mL of 20% glycerol and kept at -70 `°`C. We collectively refer to the 96 isolates collected for this study as "THRIVE-yeast" isolates. This study has been approved by the University of Manitoba Biomedical Research Ethics Board (HS24769 (B2021:026)) and Shared Health (SH2021:038).

### DNA extraction and sequencing

Genomic DNA was extracted from 24 single colonies (12 each from the vagina and rectum) from one individual with an *N. glabratus* yeast population (YST6) and three individuals with a *C. albicans* population (YST7, TVY4, TVY10). We followed a standard phenol-chloroform protocol as previously described (38). DNA quality and concentration were assessed on a Thermo Scientific™ NanoDrop 2000 and Qubit® 2.0 Fluorometer. Genomic DNA was sent to either Microbial Genome Sequencing Center ("MIGS," Pittsburgh, USA; YST6 and YST7) or SeqCoast Genomics (New Hampshire Ave., USA; TVY4 and TVY10) for sequencing. At MIGS, sample libraries were prepared using the Illumina DNA Prep kit and IDT 10bp UDI indices and sequenced on NextSeq 2000 using a 300-cycle flow cell kit, producing 2×151bp reads. The bcl-convert v3.9.3 software was used to assess read quality, demultiplex and trim adapter sequences. At SeqCoast Genomics, samples were prepared using an Illumina DNA Prep tagmentation kit and unique dual indexes. Sequencing was performed on the Illumina NextSeq2000 platform using a 300-cycle flow cell kit to produce 2×150bp paired reads. DRAGEN v3.10.11 was used to assess read quality, demultiplex and trim adapter sequences. Three isolates (TVY10R13, TVY4R4 and YST7R13) had extremely low coverage (< 20×) and were not included in genomic analysis. The average coverage from the remaining 93 isolates was 50×. The fastq files from all THRIVE-yeast have been deposited at the National Center for Biotechnology Information (NCBI) Sequence Read Archive under BioProject ID PRJNA991137.

In addition to the 93 genomes we sequenced, we downloaded an additional 182 *C. albicans* FASTQ files from the NCBI Sequence Read Archive database (39) from BioProject Accession PRJNA432884 (20) and 526 *N. glabratus* FASTQ files from 19 different projects on the SRA database (assessed on February 05, 2022), including 99 *N. glabratus* FASTQ files from SRA PRJNA361477 (40) and PRJNA669061 (41) (see Table S1 and Table S2).

### Variant calling

The sequence reads were trimmed with Trimmomatic (v0.39) (42) with standard parameters (LEADING: 10, TRAILING: 3, SLIDINGWINDOW:4:15, MINLEN: 31, TOPHRED33, following 43). Quality was assessed with FASTQC (http://www.bioinformatics.babraham.ac.uk/projects/fastqc/) and MultiQC (44). *C. albicans* trimmed paired-end reads were mapped using bwa-mem (45) to the SC5314 haplotype A reference genome (A22-s07-m01-r160) downloaded from the Candida Genome Database (46). The resulting SAM file was coordinate-sorted and converted to a Binary Alignment Map (BAM) file using samtools v1.9 (47). *N. glabratus* isolates were mapped to the CBS 138 reference genome (GCA000002545v2) downloaded from the Ensembl Genome Database (48). Alignment quality was assessed with CollectAlignmentSummaryMetrics from Picard v2.26.3 (http://broadinstitute.github.io/picard) and consolidated across all samples with MultiQC (44). All files had a >95% mapping quality. BAM files were further processed with Picard by adding a read group annotation so that samples with the same Bioproject ID had the same read group, removing duplicate PCR amplicons and fixing mate pairs. Base quality scores for the *C.albicans* aligned reads were recalibrated with known single-nucleotide polymorphisms obtained from the Candida Genome Database website (http://www.candidagenome.org/download/gff/C_albicans_SC5314/Assembly22/A22_Jones_PMID_15123810_Polymorphisms.vcf; <u>downloaded on July 29, 2020</u>) (49) using the BaseRecalibrator and ApplyBQSR from the Genome Analysis Toolkit 4.2.4.0. The average coverage for each isolate was estimated using samtools v1.9 (47).

The GATK Best Practices were adapted for variant calling. In sequence, HaplotypeCaller, CombineGVCFs, GenotypeVCFs, VariantFiltration, and SelectVariants (50–52) were used to identify single nucleotide variants (SNPs) among all sequenced isolates in diploid and haploid mode for *C. albicans* and *N. glabratus* respectively. The results SNP table was hard filtered using the suggested parameters and to match (20) (QualByDepth <L2.0, FisherStrand >L60.0, root mean square mapping quality L<L30.0, MappingQualityRankSumTest < −12.5, ReadPosRankSumTest < −8.0). We excluded variants that were called in known repetitive regions of the genome, as these are likely to reflect sequencing misalignments rather than true variants, i.e., the subtelomeric regions (15kb from the start and end of each chromosome), the centromeres, and the major repeat sequence regions, (Table S3, start and stop positions from candidagenome.org).

### Phylogeny construction

Phylogenetic trees were constructed for *C. albicans* and *N. glabratus* isolates. For each species, the multi-sample VCF file consisting of only genomic DNA SNPs was converted to a FASTA alignment using a publicly available Python script that creates an alignment matrix for phylogenetic analysis (vcf2phylip.py v2.8, downloaded from https://github.com/edgardomortiz/vcf2phylip) (53). For heterozygous SNPs in *C. albicans,* the consensus sequence is preferentially made based on the reference (haplotype A) base.

Ambiguous bases are written following IUPAC nucleotide ambiguity codes in the matrix. Ploidy is not specified, as the most parsimonious ploidy is detected by the script. The FASTA alignment was parsed in FastTree (2.1.11) (54) in the double precision mode to construct an approximate maximum-likelihood phylogenetic tree using the general time reversible model and the -gamma option to rescale the branch lengths. Compared to the time-consuming ML-based phylogeny predictors such as RAxML (55), FastTree has been found to produce large dataset trees with similar accuracy within a significantly shorter amount of time (56). The phylogeny was visualized and annotated with the Interactive Tree Of Life (iTOL, v5) (57). Isolates from *C. albicans* clade 13, a closely related but different species termed *C. africana* (58–60), were used to root the phylogeny. The *N. glabratus* phylogeny was rooted at the midpoint. While constructing the phylogeny with FastTree with this large data set yielded a phylogeny comparable to *C. albicans* whole-genome phylogenies previously generated by other groups using RAxML (20, 61), an attempt to use FastTree to resolve the relatedness of the THRIVE-yeast isolates yielded poor resolution, as most of the SNP sites were ignored during the tree construction. Hence, to generate a more refined phylogeny depicting the relatedness among the isolates from each participant, we constructed a maximum likelihood phylogeny with RAxML v8.2.12 (55) following standard practice using the GTR+G model with 20ML inferences on the alignment, inferring bootstrap replicate trees, applying MRE-based bootstrapping test, and drawing support values using TBE (Transfer Bootstrap Expectation) on the best-scoring tree.

### Pairwise SNP differences between isolates from the same site in a participant

We also looked at the variation in SNPs among isolates from the same participant within a site. Using RTG tools with the vcfsplit option (62), the VCF files for each isolate were extracted from the multi-sample VCF file. Using “bcftools query -f,” the VCF files were converted to bed files. A custom R script was used to do a pairwise comparison of the isolates from a site to determine differences in SNP positions. An ANOVA test was performed to compare pairwise SNP differences in rectal and vaginal isolates from each participant.

### *In silico* MLST typing

As clades in *N. glabratus* are commonly identified and named through MLST analysis ((40, 41)), we conducted *in silico* MLST analyses of the YST6 isolates as well as the cluster in which the YST6 isolated are associated with using stringMLST (63) which uses a predefined MLST *N. glabratus* allele library (FKS, LEU2, NMT1, TRP1, UGP1, URA3) in the PubMLST database (64). The diploid sequence types (DSTs) of the global *C. albicans* isolates were determined in a previous study using a PCR-based method (20). *In silico,* the identification of DST from the WGS data was error-prone, and known DSTs could not be replicated (data not shown).

### Determining segregating genomic blocks within isolates from the same participant

We conducted a principal component analysis of windows of genomic regions that differed among isolates from the same participant. Briefly, we used the “templated script” from the R package lostruct (local PCA/population structure, v.0.0.0.9, (65) with run parameters -t (window type): bp, -s (window size): 5000, -npc ( number of pcs): 2 and -m (number of MDS coordinates): 2. To check for possible segregation of vaginal and rectal isolates, principal component analysis plots were generated based on the extreme heterogeneous genomic windows across the entire genome.

### Relationship between average nucleotide diversity (**π**) and number of samples

We assessed the potential limitation of the sample size using average pairwise diversity differences between all possible isolate pair estimates from Pixy (v1.2.6.beta1, (66). Briefly, variants were called using GenotypeGVCFs with --all-sites option activated. Vcftools (v0.1.16) (67) was used to filter the variants (with --max-meanDP 500, --min-meanDP 20, --max-missing 0.8). Indels and mitochondrial DNA were excluded (--remove-indels, --not-chr). We compared the calculated nucleotide diversity (π) from the total number of isolates per participant (*i.e.*, *n* = 23 for TVY10, TV4, YST7 and *n* = 24 for YST6) against π estimated from a smaller number of samples (*n* = 2, 3, 4, 6, 10) from the same population. For each, we randomly selected without replacement *n* vaginal isolates 50 times, and estimated π for each sample. The mean and standard deviation were then calculated. We generated data for *n* = 12 by randomly selecting six vaginal and six rectal. We plotted π against the number of isolates included to assess the value of *n* where an asymptote for π was observed.

### Nucleotide diversity from published isolates from other studies

We compared our nucleotide diversities to instances where multiple isolates have been retrieved in either commensal or disease settings. For *C. albicans*, 3-4 isolates from 6 and 3 individuals respectively from Andersen *et al.* (21) and Sitterlé *et al*. (33) were assessed. All the isolates are from commensal settings in oral and rectal sites. For *N. glabratus*, nine to ten isolates from blood cultures in each of the nine patients (34) were compared to the 12 YST6 isolates.

### *In silico* mating-type locus detection

To determine mating type-like locus in the *C. albicans* isolates (YST7, TVY4 and TVY10), the reads were aligned to both haplotypes and consensus sequences of the MAT locus on chromosome 5 (MATa1 and MATa2 for hapA and MATα1 and MATα2) were extracted. A BLAST search was then conducted to confirm the locus. Similarly, the mating type-like locus of *N. glabratus* (YST6) isolates was determined by determining the consensus sequence for MTL1 (MTLalpha1 and MTLalpha2) and MTL3 on chromosome B, and MTL2 on chromosome E and confirming the MTL by a BLAST search.

### Loss of heterozygosity and copy number analyses

We conducted genome-wide loss of heterozygosity (LOH) and copy number variant (CNV) analyses using the web-based yeast analysis pipeline (Y_MAP_) (68). Mitochondrial DNA is by default excluded from the pipeline. Paired-end reads of the *N. glabratus* isolates were uploaded and analyzed to the CBS138 reference genome (CGD: s05-m01-r09). As *N. glabratus* is haploid, the ploidy was set as 1, which by default excludes LOH analyses. For the *C. albicans* isolates, paired-end read data for each isolate was uploaded and analyzed against the SC5314 A22-s02-m09-r10 reference genome. Ploidy was left at the default value (two, i.e. diploid), and correction was enabled for GC-content bias and chromosome-end bias. THRIVE-yeast isolates were compared against the SC5314 haplotype map to distinguish between ancestral and newly evolved LOH signatures and determine which allele was retained. CNV profiles were compared to the reference SC5134 isolate as well as closely related isolates in the phylogeny. The genomic elements within observed CNV regions were identified using the "gene/sequence resources" section of the Candida Genome Database (CGD, https://candidagenome.org).

### Growth Rate Assay

Two separate growth rate assays were conducted to measure growth in RPMI (10.4 % w/v RPMI powder, 1.5 % w/v dextrose, 1.73 % w/v 3-(N-morpholino) propanesulfonic acid (MOPS), adjusted to pH 7 with NaOH tablets) and vaginal simulative medium ("VSM", following (20): 1.16 % v/v 5 mM NaCl, 3.6 % v/v 0.5 M KOH, 0.0128 % v/v 99% glycerol, 20 % v/v 0.01 M Ca(OH)_2_, 1.34 % v/v 0.5 M Urea, 6.6 % v/v 0.5 M glucose, 0.67% w/v solid YNB, 0.85 % v/v 2 M acetic acid, 0.192 % v/v lactic acid, adjusted to pH 4.2 with NaOH tablets). For each experiment, 5 µL of frozen glycerol stock from all THRIVE-yeast isolates was inoculated in duplicate into 500 µL of RPMI or VSM and incubated for 48 h at 37 °C with agitation at 250 rpm. Cultures were then standardized to an optical density (OD) of 0.01 A600 in RPMI or VSM, and 200 µL was transferred into a 96-well round bottom plate and sealed with a Breathe-Easier sealing membrane (Electron Microscopy Sciences, PA, United States). OD_600_ readings were taken by the Epoch plate reader (Biotek) every 15 minutes, with continuous shaking at 37 °C for 48 h.

From each well, the maximal growth rate was calculated as the spline with the highest slope using a custom R script written by Dr. Richard Fitzjohn (https://github.com/acgerstein/THRIVE-variation/scripts_real/growthrate_generic.R). The average growth rate between two technical replicates for each isolate in each growth medium was used for visualization and statistical analysis. Statistical outliers were determined through Rosner’s test of outliers available through the rosterTest function in the EnvStats R package (69). For each population, we started with a k value of one (i.e., testing for a single outlier). If that was significant, we increased k by one until no additional outliers were identified.

### Drug resistance and tolerance

Disk diffusion assays were carried out to measure resistance and tolerance. A pilot experiment was done on 24 isolates from YST7 in five different drugs (FLC: fluconazole, CLT: clotrimazole, MCZ: miconazole, NYT: nystatin, BA: boric acid). Subsequently, disk diffusion assays were conducted on all isolates to fluconazole and boric acid at pH 4.2. We chose to focus our efforts on fluconazole and boric acid as these are drugs in different classes that are both treatment options for induction and maintenance therapy of recurrent VVC and boric acid is used in treatment of non-albicans VVC (van Schalkwyk et al. 2015). Standard CLSI M44 guidelines for antifungal disk diffusion susceptibility testing (70) were generally followed except for pH adjustment as appropriate. To initiate each experiment, frozen stock of all THRIVE-yeast was streaked onto Sabouraud dextrose agar (SDA) plates and incubated for 72 h at room temperature. Isolates were then subcultured by streaking one colony onto a fresh SDA plate and incubating at 37 °C for 24 h. Colonies from each isolate were then suspended in 200 μL of 0.85% saline solution and standardized to an OD_600_ of 0.01 in 1 mL of saline solution. Within 15 minutes of standardization, 100 μL of the culture was spread evenly using autoclaved glass beads on Mueller-Hinton (MH) plates that had been adjusted to a pH of 4.2. Plates were left to dry for 20 minutes before placing a single 5 mg boric acid or 25 mg fluconazole disk in the center of the plate. Plates were incubated upside down at 37 °C. After 48 h, photographs of each plate were taken on a lightbox. Previous work demonstrated consistent drug resistance values at 24 h and 48 h. The entire experiment was conducted twice for each isolate × drug × pH combination, with two technical replicates for each of the two biological replicates.

The 48 h images were processed in ImageJ as previously described (71). They were cropped to a uniform size, the colours were inverted, and brightness and contrast were adjusted to maximize the contrast between the white disk and the black background. The adjusted images were run through the diskImageR package (72) for drug resistance (RAD_20_) and tolerance (FOG_20_) quantification. Briefly, diskImageR calculates resistance as RAD_20_, as the radius of the zone of inhibition where growth is reduced by 20% relative to growth on the margins of the plate where there is no drug, and tolerance as FoG_20_, the fraction of realized growth between RAD_20_ and the disk.

A Welch two-sample t-test that did not assume equal variance was used for each participant × drug combination to compare vaginal and rectal isolates. Statistical outliers were determined through Rosner’s test of outliers for growth rates. All statistical analysis was done at a type I error rate of 0.05.

### Invasive growth assay

To examine invasive growth, we revised the methods from (73). Freezer stock from THRIVE-yeast isolates were streaked onto 20 mL yeast peptone dextrose (YPD) plates (2% w/v peptone, 2% w/v yeast extract, 1.8% w/v agar, 1% w/v glucose, 0.00016% w/v adenine sulphate, 0.00008% w/v uridine, 0.1% v/v of chloramphenicol and ampicillin) and grown for 72 h at room temperature. A single colony was then randomly chosen from each isolate and inoculated into 200 μL YPD. If no single colonies were available, a similar amount of culture from the colony lawn was used. Cultures were standardized to OD_600_ 0.01 in 1 mL of liquid YPD media, then 2 μL of standardized culture was spotted onto the surface of a 20 mL solid YPD plate in a hexagonal pattern for a total of 7 spots per plate (i.e., spotted at each vertex and in the center). Plates were incubated for 96 h at 37 °C. The surface growth was washed off using distilled water and a photograph was taken in a dark room on a lightbox. Two biological replicates were performed for each isolate.

The qualitative amount of invasive growth for each isolate was determined by visual examination of the post-wash photographs. To develop a five-point scale, two different people independently went through the post-wash pictures from YST6 and YST7 and selected two to six representative pictures that fit into five levels of invasive growth (scored as 1-5). The independent selections were then compared, and one image from each person was chosen as the most representative for each level of the scale. Using these as a reference, each isolate was then categorized into the five levels of the scale (0 - no growth/pipette tip indent, 0.25 - pinprick growth, 0.5 - circular growth evident, 0.75 - circular growth with pinprick, 1 - dense growth throughout). The maximum score between the two bio-replicates of each isolate was used for statistical analysis, though the same statistical conclusions were obtained if the mean score was used instead. For each participant, a Wilcoxon rank sum test was used to compare vaginal and rectal isolates.

## Results

### THRIVE-yeast isolates in the global species phylogenies

Participants with a history of RVVC were recruited from a specialty yeast clinic in Winnipeg, Canada. Vaginal and rectal isolates were collected from swabs plated onto SDA and chromogenic *Candida* agar from seventeen participants who were enrolled and screened intermittently between January 2020 and November 2022. As the goal was to quantify standing genetic variation from single time point vaginal and rectal populations, we haphazardly isolated vaginal and rectal yeast isolates from each of the four participants where we had at minimum 12 isolates from each site on our selective medium plates, for a total of 96 isolates which we refer to collectively as the “THRIVE-yeast” isolates. Isolates from one participant were *N. glabratus* (YST6), while *C. albicans* was isolated from the other three (TVY4, TVY10, and YST7). Three isolates (TVY10R13, TVY4R4 and YST7R13) had low depth of coverage (< 20×) and were excluded from the genomic (but not phenotypic) analyses. All *C. albicans* isolates were MAT-heterozygous diploids (a/α), while the *N. glabratus* isolates were all MTL1a.

The phylogenetic relationship of the THRIVE-yeast isolates was evaluated in the context of available short-read whole-genome sequenced (WGS) isolates from each species. We exhaustively searched NCBI for available *N. glabratus* sequences, finding and downloading fastq data from 526 isolates (Table S1). There was an over-representation of blood isolates (n = 430, > 80%), and 53 isolates did not have a listed isolation site. Notably, we only found a single vaginal isolate and twelve stool isolates. Additional isolation sites (e.g., abdomen, peritoneal fluid, mouth, catheter, bronchioalveolar lavage, urinary tract) were similarly represented by very few isolates. This distribution precludes an opportunity to determine whether or how *N. glabratus* site of isolation contributes to isolate relatedness. We added the fastq data from the YST6 isolates to the 526 global isolates and used this data to construct the largest *N. glabratus* phylogenetic tree to date.

The YST6 isolates are monophyletic and cluster with 68 bloodstream isolates and three isolates of unknown provenance from the United States, Canada, and Australia (Fig. 1A). Unfortunately, over 80% of the sequenced isolates are bloodstream infection isolates, with only one previous isolate annotated as originating from the vagina, rectum, or stool (Table S1), precluding further analysis. The globally published *C. albicans* phylogeny comprises 182 isolates from a wide breadth of geographic and anatomical sites (20). Isolates from TVY4, TVY10 and YST7 all form monophyletic groups and fall within a subgroup that contains twenty-three additional isolates in clade 1, the most common clade (Fig. 1B). TVY4 and TVY10 isolates are beside each other and shared a common ancestry with M40 which is also a vaginal isolate from Morocco. The YST7 isolates are most closely related to three vaginal isolates (one each from Brazil, Morocco, and China) and one oral isolate from Niger. Seven of the remaining eighteen isolates in the clade 1 subgroup were also isolated from the vagina. This is a statistical enrichment for vaginal isolates compared to the rest of the isolates in clade 1 (THRIVE-yeast isolates from each participant were counted as a single isolate; Fisher exact test comparing 14 vaginal isolates out of 26 in the subgroup to 3 vaginal isolates out of 17 in the rest of clade 1, p = 0.026). If we discount the 35 predominantly vaginal isolates in clade 13, which is now recognized as likely a separate species (*C. africana)* (58, 60), clade 1 as a whole is also statistically overrepresented for vaginal isolates compared to the *C. albicans* tree in general (17 vaginal isolates in clade 1 out of 43 total isolates, compared to 18 vaginal isolates out of 107 total isolates; Fisher exact test, p = 0.005). Thus, although the sequenced vaginal isolates are located in six different clades, they are over-represented in clade 1 relative to a neutral expectation that vaginal isolates are equally likely to be found anywhere in the existing tree.

**Fig. 1.**
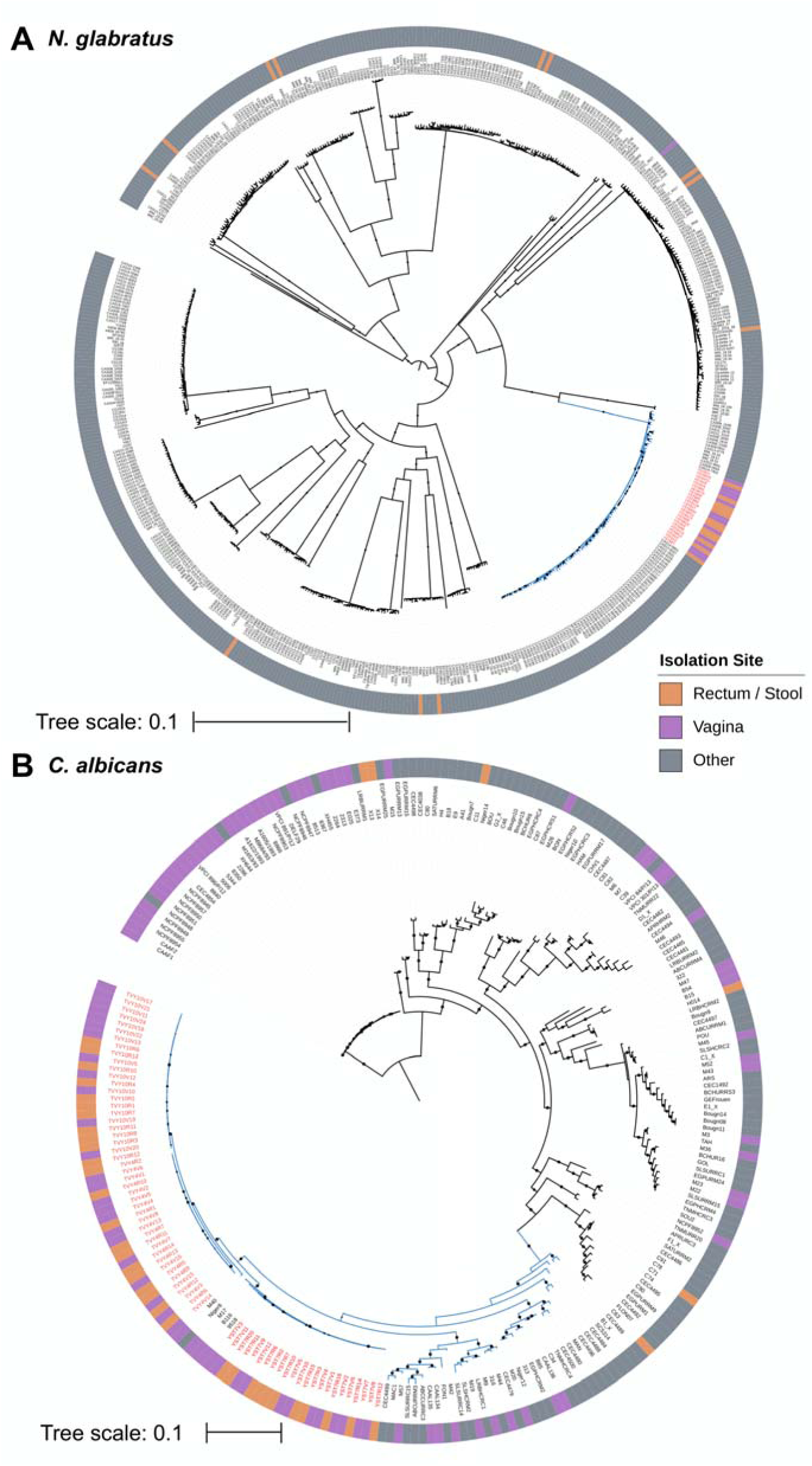
Approximate maximum likelihood phylogenies of (A) *N. glabratus*, including 526 global isolates, and YST6 vaginal and rectal isolates, and (B) *C. albicans*, including 182 isolates from Ropars et al. 2018 and vaginal and rectal isolates from TVY4, TVY10 and YST7. THRIVE-yeast isolates are indicated with red labels. The YST6 *N. glabratus* isolates are in a cluster of ST16 isolates, while the TVY4, TVY10, and YST7 *C. albicans* isolates are in clade 1 (blue branches indicate A) ST16 and B) clade 1 isolates). The *N. glabratus* phylogeny was rooted at the midpoint, and the *C. albicans* tree was rooted by *C. africana* isolates (grey labels).

The *N. glabratus* phylogeny is composed of long internal branches with only a small amount of variation within the clades; clades defined through MLST are mainly consistent with WGS (40, 41). An *in silico* MLST analysis of our WGS data found that the YST6 isolates cluster with 71 isolates: 67 ST16, two ST60, and two ST187. ST60 is distinguished by a single polymorphism at position 298 in *LEU2,* while ST60 is distinguished by a single polymorphism at position 250 in *NMT1*. *C. albicans* diploid sequence types (DST) do not map as well to WGS clades. Each of the clade 1 subgroup isolates has a different DST (20), and the differences are mainly registered in 39 positions across the genes.

### Vaginal and rectal isolates are closely related and phylogenetically overlapping

Following the observed monophyly among isolates from the same participant, we next assessed the relatedness of vaginal and rectal isolates. We first generated phylogenies for each individual using RAxML (55), as previously done for closely related bacteria strains (74). The vaginal and rectal isolates from all four participants are phylogenetically overlapping (Fig. 2A). Few branches within the four phylogenies had bootstrap support exceeding 80%, yet well-supported clusters in all participants included isolates from both body sites. For each population, we also conducted a local principal component analysis (PCA) that examines the regions of high genomic heterogeneity within populations. From all participants, the regions of high differences were generally distributed throughout the genome, and the PCA failed to segregate vaginal and rectal isolates (Fig. S1).

**Fig. 2.**
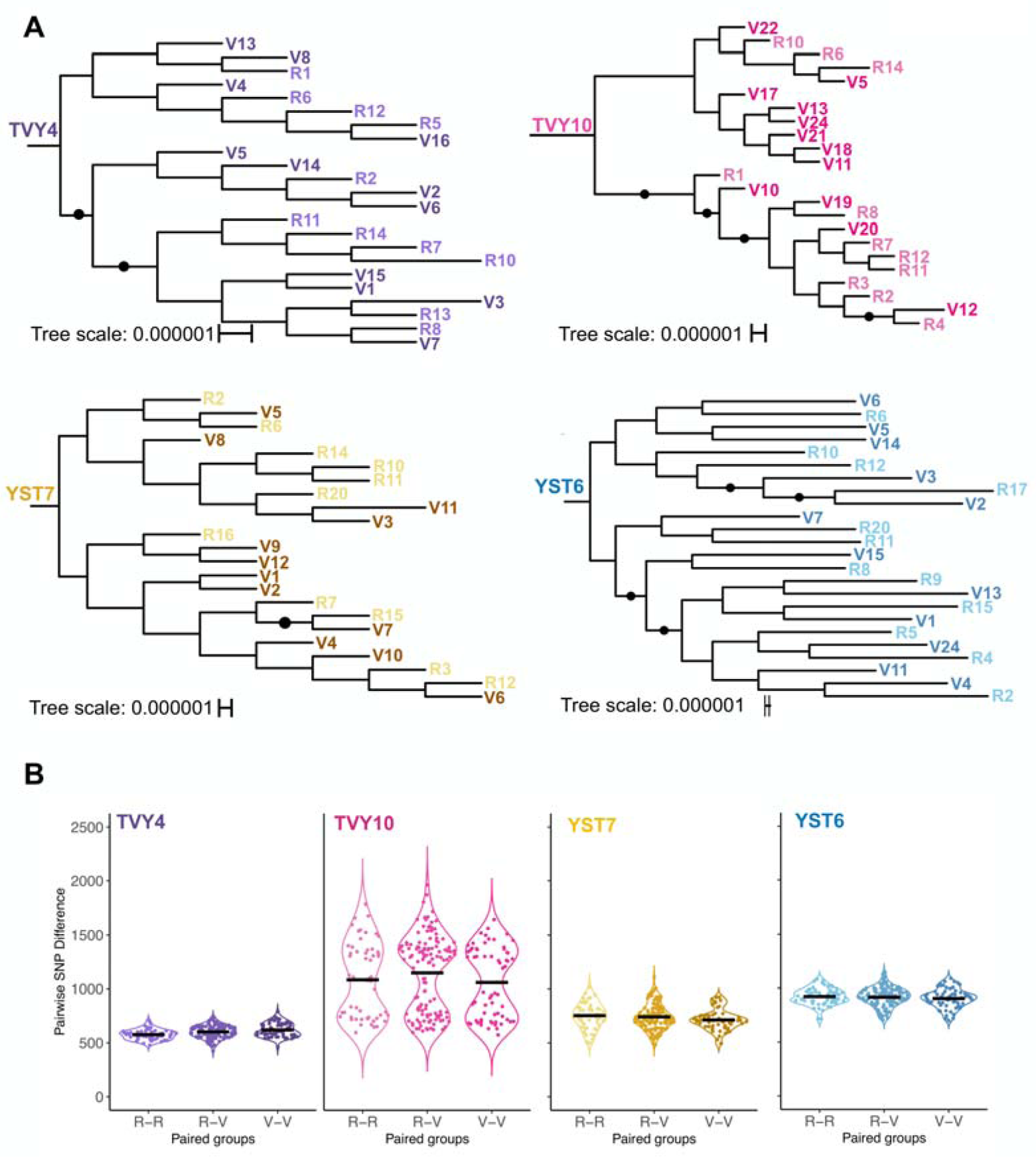
Within participant phylogenetic and single nucleotide position (SNP) analyses. (A) The fine-scale phylogenetic structure among THRIVE-yeast isolates. Vaginal (V) and rectal (R) isolates were acquired from four participants with a history of RVVC (YST6: *N. glabratus*; TVY4, TVY10, YST7: *C. albicans*). The isolate numbers are arbitrary based on the order they were collected off culture plates. Reads were aligned to their respective reference genomes (SC5314 and CBS 138), and variants were called using a custom variant calling pipeline adopted from the GATK best practices. Mitochondrial variants, as well as variants in repeat regions, were excluded. Phylogenetic trees were generated with RAxML; black circles indicate branches with bootstrap support ≥ 0.8. (B) Within-participate pairwise comparison of single nucleotide positions between isolates from the indicated sites (V-V: vaginal-vaginal, R-V: rectal-vaginal, R-R: rectal-rectal).

### Pairwise differences in single nucleotide polymorphisms among isolates per participant

Although isolates were closely related, WGS data picked up SNP variation among all pairs of participant isolates. To test whether within-population vaginal diversity was lower than within-population rectal diversity, we compared pairwise SNP differences among vaginal isolates to the pairwise SNP differences among rectal isolates. To test whether there was a signal of divergence between sites, we also compared single-site differences to pairwise differences between isolates across sites. The average pairwise SNP differences between vaginal isolates were very similar to average pairwise SNP differences between rectal isolates and between isolates from different sites (Fig. 2B). The only significant difference was in TVY4, where the average SNP differences among rectal isolates were significantly lower than the vaginal isolates (ANOVA test, YST7: F_2, 250_ = 2.164, P = 0.117; YST6: F_2, 250_ = 0.785, P =0.457; TVY4: F_2, 250_ = 8.599, p =0.000244; TVY10: F_2, 250_ = 1.66, P = 0.192; Tukey’s HSD Test for multiple comparisons; p = 0.0001). The distributions were fairly normal, as expected for isolates with low population structure, except that TVY10 isolates showed a bimodal distribution of pairwise SNP differences in both the vaginal and rectal sites.

### Minimum number of isolates for estimating standing genetic variation within participants

A major goal of our work was to compare diversity within RVVC populations to diversity observed in other contexts to make inferences about the evolutionary process based on the observed degree of standing genetic variation. However, the small number of comparable studies all sequenced different numbers of isolates, and nucleotide diversity (π) will decrease with an increased number of samples taken from a population. To quantify the scale of this effect of changing the number of isolates, we conducted a bootstrap analysis using our 12 vaginal isolates from each individual. We repeatedly resampled 3-10 isolates and recalculated diversity. There was subtle variation in nucleotide diversity among the three participants with *C. albicans* populations. In all cases, the shape of the diversity curve with the number of isolates was very similar---an elbow was observed around n = 6 (Fig. 3). Nucleotide Diversity in YST6 (*N. glabratus*) was two orders of magnitude lower, and there was not a consistent change in diversity with the number of isolates.

**Fig. 3.**
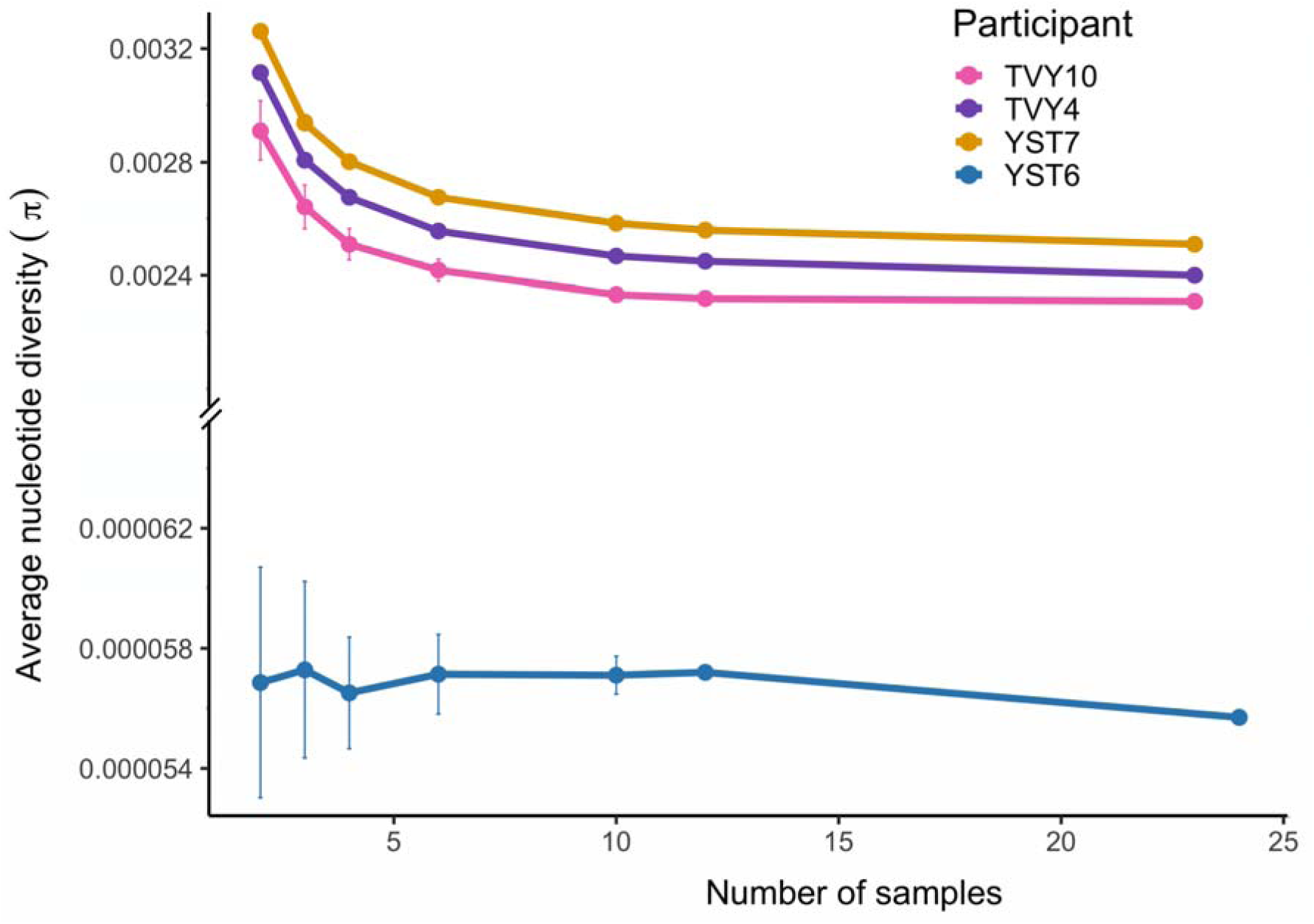
Relationship between sample size and average nucleotide diversity. π was calculated for different numbers of samples from each individual. For n= 2, 3, 4, 6, or 10, the given number of samples was randomly selected from the vaginal isolate set. For n = 12, the datasets were generated by randomly choosing 6 samples from each of the rectal and vaginal isolates sets. The bootstrap analysis was done 50 times for each sample size, and the mean and standard deviation among data sets were calculated. For n = 23 (*C. albicans*, TVY10, TVY4, YST7) or 24 (*N. glabratus,* YST6), the nucleotide diversity of all samples was calculated.

### THRIVE-yeast isolates share similar diversity as isolates from commensal and other disease settings

We downloaded the fastq files from two previous *C. albicans* studies on commensal populations (21, 33) and used our pipeline to calculate the average nucleotide diversity for each. For all populations, including our own, where necessary we down-sampled the number of isolates to three, consistent with the lowest number of isolates sampled from the commensal populations. The average nucleotide diversity was very similar across most populations (Fig. 4A). The exception was populations from two participants previously shown to have isolates from different phylogenetic clades. The YST6 vaginal isolate population was compared to reanalyzed fastq data from nine different BSI populations from Badrane et al., which all had between 9-10 isolates (34). The average nucleotide diversity from our population was similar to the diversity from four participants yet much higher than the average nucleotide diversity from the other five (Fig. 4B).

**Fig. 4.**
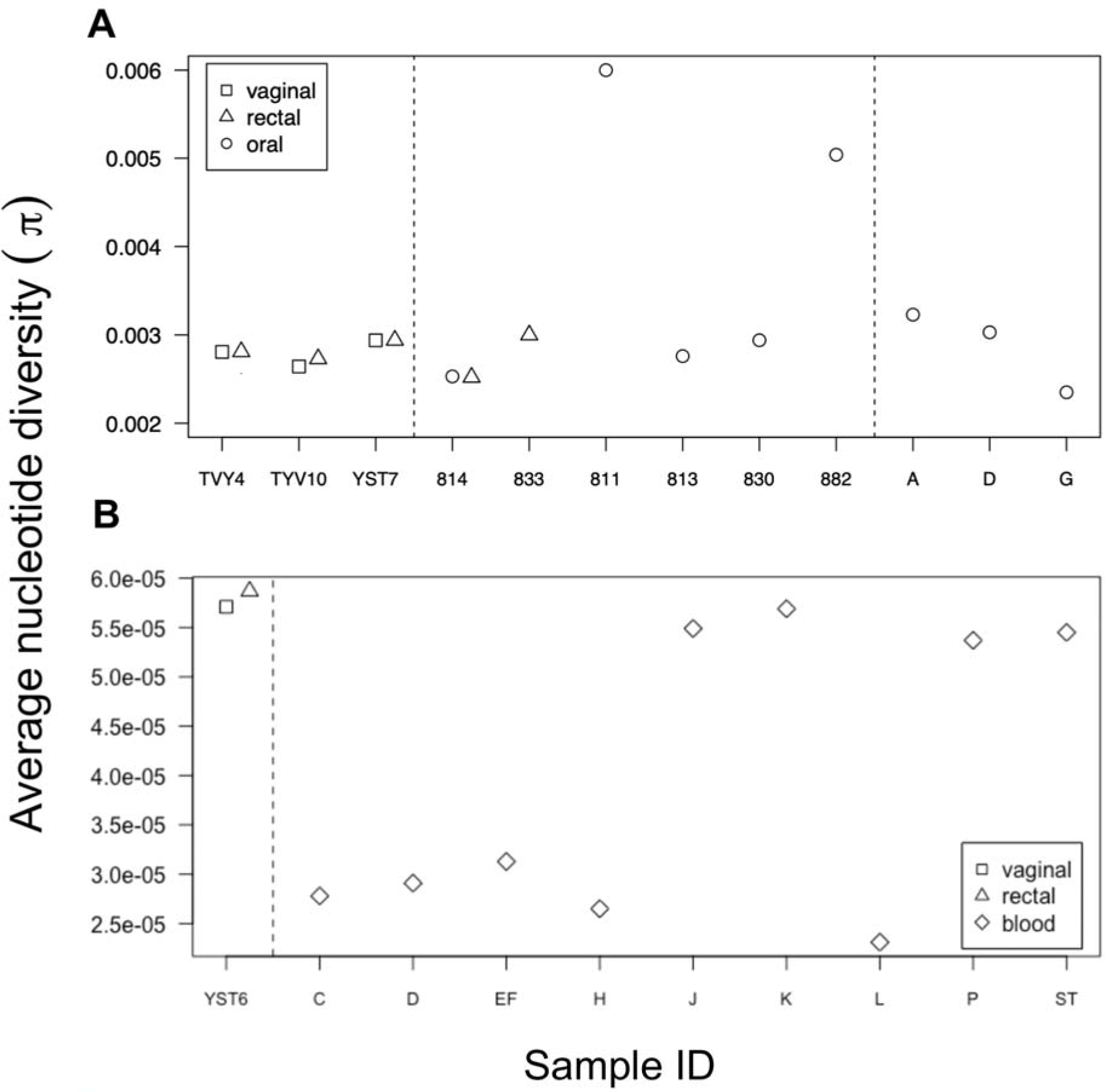
Comparison of the average nucleotide diversity between our samples and isolates from other studies. (A) Comparison of π in THRIVE *C. albicans* isolates from TVY4, TVY10, and YST7 to commensal isolates from two previously published studies. For accurate comparison, three randomly chosen samples were subsampled from each site in the THRIVE-yeast isolates. (B) Comparison of π in THRIVE *N. glabratus* (YST6) isolates to bloodstream infection isolates from 10 individuals in a previous study.

### Little variation in copy number or loss of heterozygosity within populations

We examined copy number variation (CNV) and loss-of-heterozygosity (LOH) events among THRIVE-yeast isolates and their closest relatives using Y_MAP_ (68). No CNVs were identified in any YST6 isolates (Fig. 5A). A single ∼50 kb CNV on the right arm of chr3 was identified in all YST7 isolates (Fig. 5B). This CNV is also present in the closest relative to the YST7 isolates, vaginal strain 9518, but is absent in the next two closely related strains that are also vaginal in origin (B116 and M17). As Y_MAP_ visualizations are based on averages across 5000 bp sliding windows, we examined the region in finer detail. Coverage was measured from the BAM files to compute the depth at each position in that region. Mapped coverage was inconsistent with the profile of a typical CNV; the majority of the region had only a slightly elevated copy number relative to the rest of the genome (Fig. 5C). Two small (< 200 bp) regions spiked up to ∼6-fold and ∼14-fold coverage, the first internal to *ALS6* and the second to *ALS7* (Fig. 5C). A third region comprised of elevated coverage maps to another gene with close homology to other genes in the genome, *CYP5*, a putative peptidyl-prolyl cis-trans isomerase (75). The region identified in Y_MAP_ is thus likely to primarily reflect an error in mapping rather than a true CNV with potential biological effects. No other CNVs or aneuploidies were identified in the other THRIVE-yeast isolates.

**Fig. 5.**
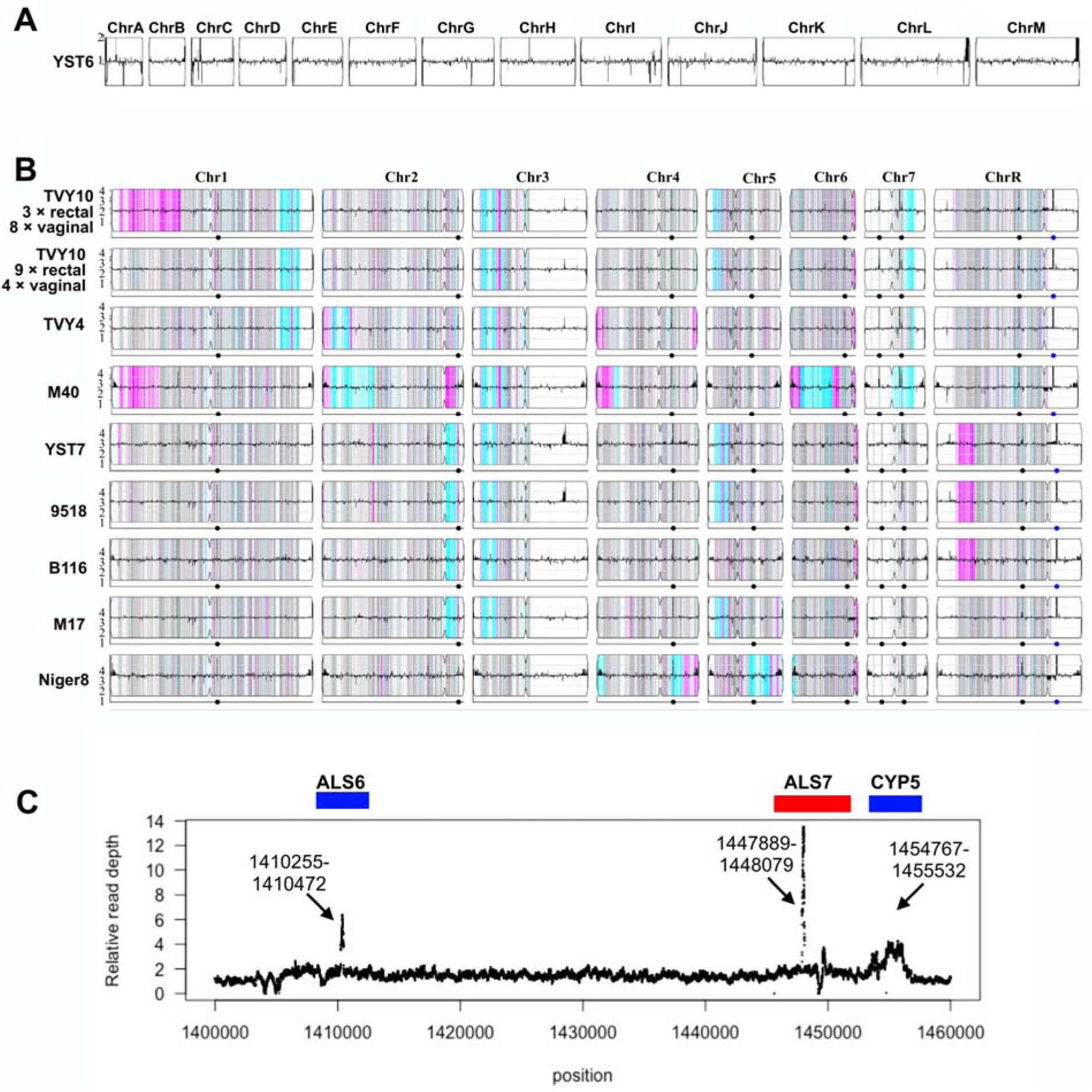
CNV and LOH profiles of THRIVE-yeast. (A) Representative trace of a YST6 isolate (*N. glabratus*) and (B) TVY10 isolates, representative traces of TVY4 and YST7 isolates with the closely related M40 isolates to them, and the four isolates that are closely related to YST7 (three vaginal isolates and Niger8, oral isolate). The relative copy number compared to the reference genomes is shown as the horizontal black line with the scale on the y-axis. For *C. albicans* isolates, the density of heterozygous SNPs is shown as the vertical lines spanning the height of each chromosome box, with the intensity representing the number of SNPs in 5 kb bins. Heterozygous SNPs are gray, homozygous SNPs are colored based on the retained SC5314 homolog: cyan for “AA” and magenta for “BB.” White indicates an ancestral LOH in SC5314. For each chromosome, the centromere is indicated by an indentation in the box. The dots on the bottom line below each box indicate the positions of major repeat sequences. (C) Fine-scale coverage mapping of the putative CNV on chr3. Shown is one representative trace from YST7 R2, all isolates have a similar pattern. Gene positions above the figure are approximated.

LOH analysis was consistent with the phylogenetic analysis and diversity metrics. Generally, all isolates from the same participant shared an LOH profile. Among the THRIVE-yeast *C. albicans* isolates compared to the SC5314 reference, nearly all chromosomes had at least one LOH event extending to the telomere (Fig. 5B, Fig. S2). No large interstitial LOH events were identified. TVY4 isolates share a complicated LOH region on the left arm of chr2 that flips back and forth from the two haplotypes, as well as small LOH regions on both the left and right arms of chr4. Both TVY4 and TVY10 isolates have unique LOH regions in common with their closely related isolate (M40) which is also a vaginal. However their LOH regions on M40 are of varying sizes compared to their counterpart regions in TVY4 or TVY10. YST7 shares LOH blocks on the right arm of chr2 from the centromere to the telomere, on the left arm of chr5, and on the left arm of chrR with the vaginal isolates it is most closely related to (i.e., 9518, B116, M17) but not Niger8, an oral isolate. The exception of wholly shared LOH blocks among participant isolates was in TVY10, where a single LOH event on the left arm of chr1 was present in twelve TVY10 isolates that cluster together on the phylogeny.

In addition to the characteristic LOH regions within populations, some LOH regions were also shared between isolates from different participants. All isolates have an LOH block on the left arm of chr3 and the right arm of chr6. TVY10 and TVY4 isolates share a ∼300 kb region on the right arm of chr1. A LOH region on chr7 beside the ancestral chr7 right arm LOH block is also present in both TVY10 and TVY4, though it is larger in TVY10. TVY10 and YST7 (but not TVY4) both have LOH blocks on the left arm of chr5, though the allelic profile between them is different.

### Phenotypic variation

We quantified within-population phenotypic variation in parallel to genotypic variation. The average growth rate for YST6 *N. glabratus* isolates was higher than the *C. albicans* populations in both Roswell Park Memorial Institute (RPMI) medium (Fig. 6A) and vaginal simulative medium (VSM, Fig. 6B). Growth rates were either the same between vaginal and rectal isolates or the rectal isolates were higher when grown in either medium (Welch Two Sample t-test, Table S4, rectal isolates higher in TVY4 grown in RPMI: t = 2.50, df = 16.1, p-value = 0.023; TVY4 grown in VSM: t = -2.36, df = 18.1, p-value = 0.030; YST7 grown in VSM: t = -3.63, df = 18.8, p-value = 0.002). Consistent with a visual inspection, single statistical rectal outliers with increased growth rate relative to other isolates were seen in YST7 and TVY4 grown in RPMI, and YST6 and TVY10 grown in VSM (Fig. 6, Rosner’s test for outliers). No additional outliers were identified in the other populations.

**Fig. 6.**
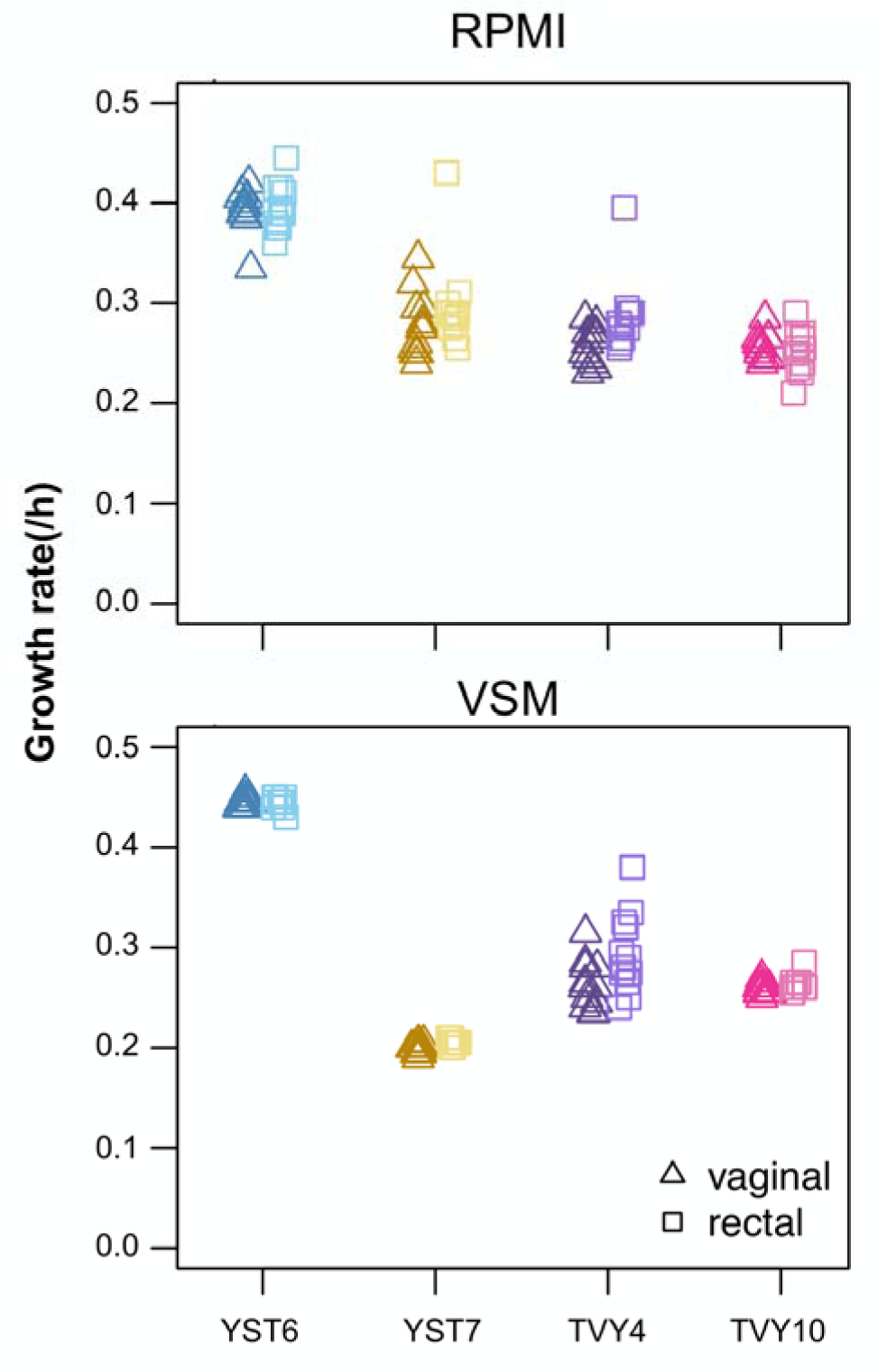
Growth rate was measured from 12 vaginal and 12 rectal isolates from each population. Optical density was recorded every 15 minutes in a plate reader with constant shaking and incubation at 37 ℃. Each point represents the mean of two technical replicates for each of the two biological isolates, 24 isolates were measured for each group. The growth rate was calculated as the spline with the highest slope using a custom R script.

We then compared drug resistance and drug tolerance from vaginal and rectal isolates. We conducted a pilot experiment on 24 isolates from participant YST7 to quantify variation in drug resistance for five different drugs that are indicated as treatment options by the Society of Obstetricians and Gynaecologists of Canada for uncomplicated, recurrent and non-albicans VVC (van Schalkwyk et al. 2015). We also examined drug tolerance, the ability of drug-susceptible populations to grow slowly in the presence of high levels of fungistatic drugs, which also emerged as a trait that varies among different fungal species and isolates [25,26].

Tolerance may be implicated in the propensity to cause fungal disease in other contexts [27] but has not previously been examined in the context of R/VVC. We found very little variation among the isolates for either phenotype in any drug (Figure S3). Given that, we proceeded with quantifying drug responses for just fluconazole and boric acid at pH 4.2, as these are drugs from different classes that are commonly prescribed in our local clinic. The site of isolation was only significant for YST6 BA resistance (vaginal isolates were slightly more tolerant than rectal isolates; t-test, t = 2.77, df = 18.2, p-value = 0.012, Table S4, Fig. 7). A visual examination of the data also indicated that the majority of isolates were very similar to each other. Formal outlier statistical tests that grouped vaginal and rectal isolates were broadly consistent with the qualitative visual assessment, identifying only two outlier isolates for tolerance (both rectal isolates in BA, one from YST6, one from YST).

**Fig. 7.**
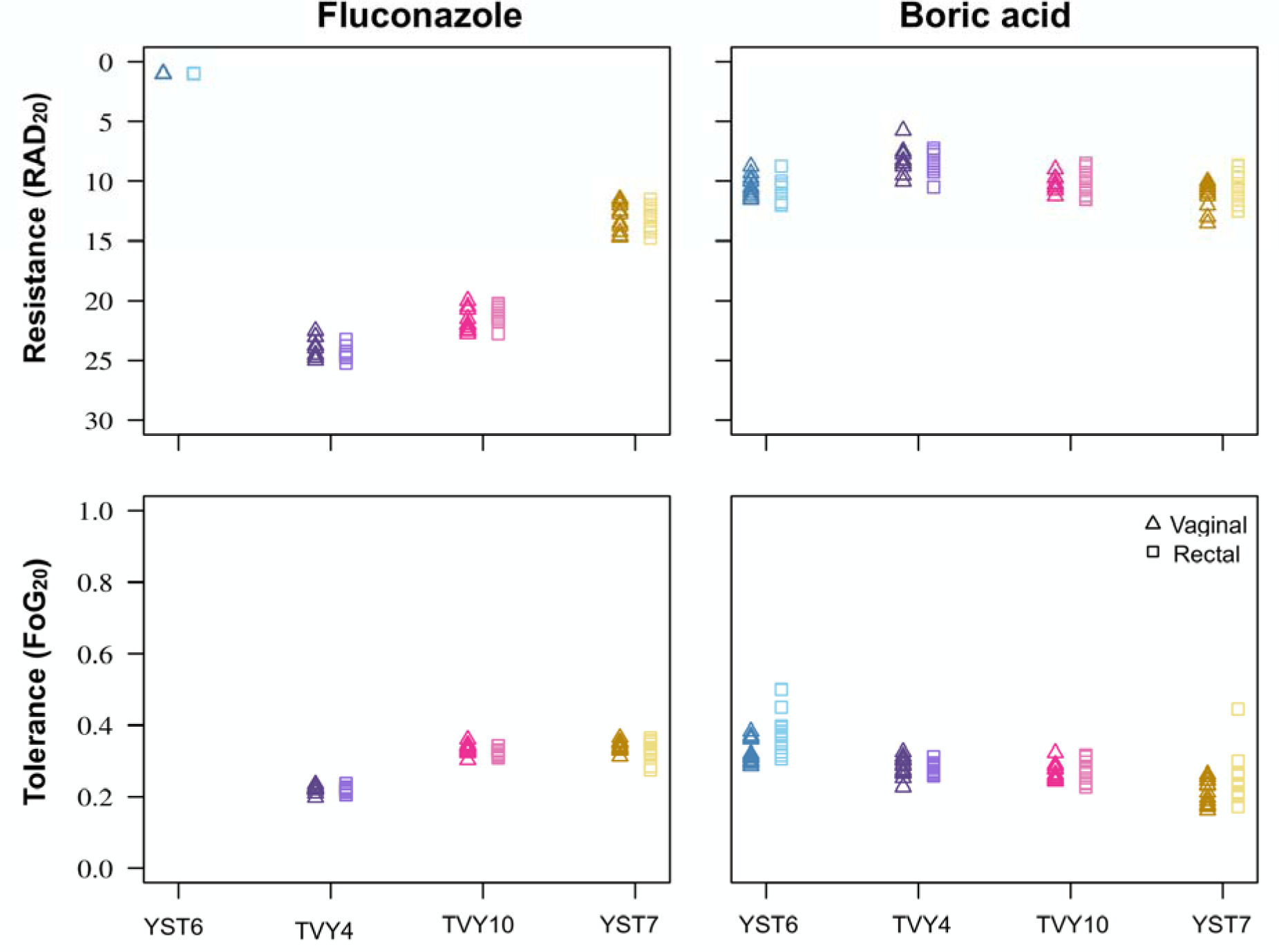
Drug response phenotypes from disk diffusion assays. Drug resistance (top panels) and drug tolerance (bottom panels) were measured from disk diffusion assays for fluconazole (FLC) and boric acid (BA). Drug response was measured on pH 4 Mueller-Hinton plates using the R package *diskImageR* (72), which computationally measures response parameters from images of disk diffusion assays. Each point represents the mean of four replicates (two technical replicates for two biological replicates).

There was considerable variation among participants and isolates for invasive growth (Fig. 8, Table S4). However, there was no difference between YST6 or TVY10 vaginal and rectal isolates. In contrast, YST7 vaginal isolates had higher invasive growth than rectal isolates (W = 115.5, *P* = 0.0002), and TVY4 rectal isolates exhibited higher invasive growth than vaginal isolates (W = 372, p-value = 0.032). The overall picture is thus that invasive growth seems to vary more between participants than between sites of isolation and that statistical differences between sites are likely due to neutral processes rather than selection for invasive growth in the vaginal environment.

**Fig. 8.**
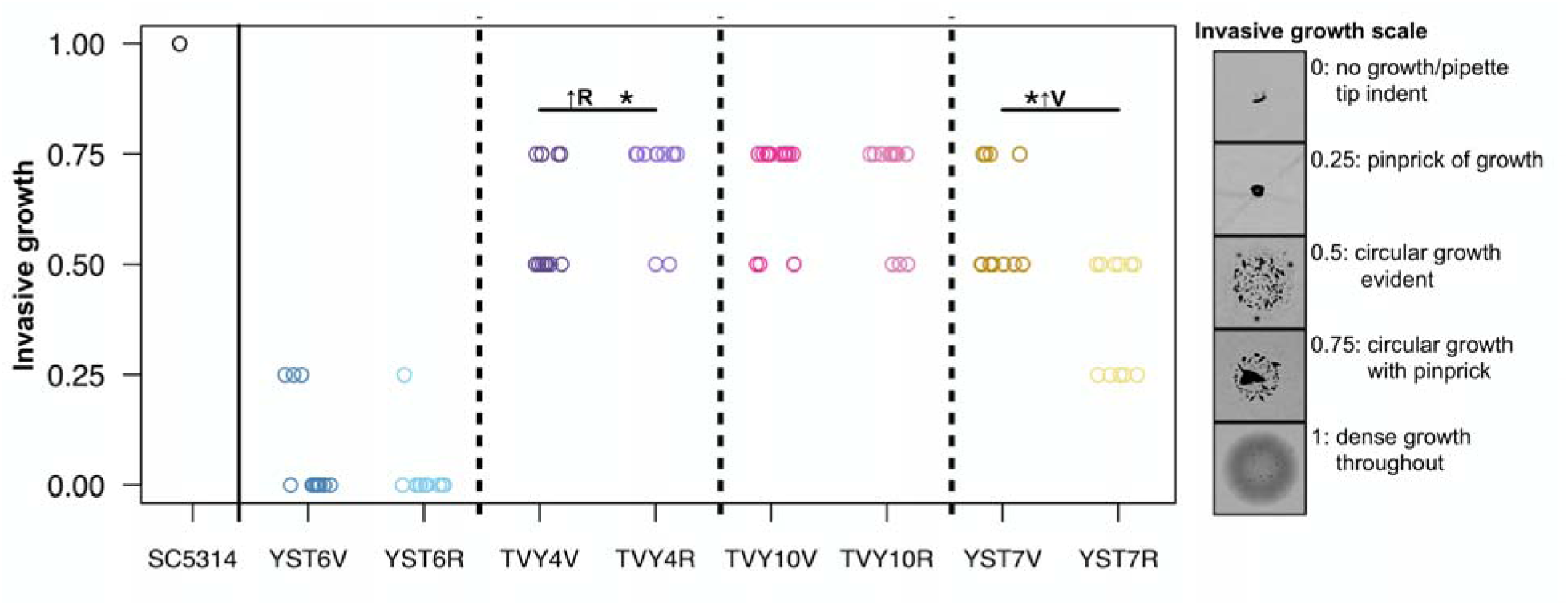
Invasive growth was qualitatively scored after growth on YPD plates for 96 h. Each point indicates the maximum score between two bio-replicates for each isolate.

## Discussion

The biological basis of RVVC needs to be better understood. To study the vaginal yeast populations implicated in the disease, we quantified the diversity of 12 vaginal and 12 rectal isolates from four people with a history of RVVC who had large yeast populations at both sites at the time of sampling. In each case, the isolates formed monophyletic groups, and the vaginal and rectal isolates were phylogenetically overlapping, consistent with a common ancestral source and frequent migration between the two sites. This is in concordance with previous genetic studies that used more coarse sequencing methods, which found high genetic similarity between vaginal and rectal isolates in women with R/VVC (76–78). Our phenotypic analyses are also consistent with the genetic results; there was minimal diversity in drug responses or growth ability in clinically relevant medium and no consistent difference between isolates from different isolation sites for invasive growth. Previous studies have also found that virulence factor phenotypes have similar expression among vaginal and rectal isolates in the context of R/VVC (79) and among oral and rectal isolates from healthy individuals (21). We thus found no evidence for selection acting differently at the two sites at either the genotypic or phenotypic levels.

Multiple potential evolutionary explanations are consistent with our results. It could be that the selective pressures that most influence adaptation are similar in both environments, leading to selection for the same traits. However, it could also be that selection cannot overcome either (or both) migration or genetic drift due to a low effective population size. The isolates come from individuals with a long history of repeatedly taking antifungal drugs, and the yeast population sizes likely undergo many bottleneck cycles over time, functionally reducing the efficacy of selection. Nevertheless, we did observe 100s of SNP differences between all isolate pairs, indicating the presence of what has been termed microvariation, and could be of the magnitude sufficient for adaptation. These results underscore the need for further investigations to understand the relationship between populations at different body sites and over time in individuals with RVVC.

The observed average nucleotide diversity among populations for the RVVC isolates was higher than anticipated. *A priori,* we had predicted that genetic diversity would be highest in commensal populations from healthy individuals at sites that do not have obviously strong selective pressures acting on them (i.e., they are from common commensal sites from individuals that are not regularly taking drugs) and lowest in bloodstream infections which are known to have a small population of circulating yeast cells. However, genetic diversity within the three RVVC *C. albicans* populations was similar to what we calculated from oral and rectal isolates from seven healthy individuals (though lower than two commensal oral populations previously known to have isolates from different clades, (21, 33). The only appropriate *N. glabratus* study we could find involved nine individuals with bloodstream infections. The RVVC *N. glabratus* population had similar diversity as the four BSI populations from an ST group (ST3) that is closely related to ST16, the cluster YST6 isolates are in, yet was much higher than the other five patient isolate populations from more distantly related clades (34). Future work in *N. glabratus* will more deeply explore whether there is a consistent relationship between clade and within-population genetic diversity. Given the limited sample size available for different contexts, it is difficult to make inferences about the evolutionary dynamics occurring in these different contexts. There may be selection in the commensal oral (80) and rectal environments, involving the fixation of alleles and reducing genetic diversity. It could also be that the population bottlenecks in RVVC (and in at least some bloodstream infections) are not as strong as anticipated. Teasing apart these explanations will require data on many more yeast populations. As whole genome sequencing becomes ever more routine, we hope additional studies will examine intra-population diversity, as we lack sufficient benchmarks to properly contextualize whether diversity is truly “high” or “low” relative to other contexts.

Importantly, we chose isolates for sequencing blind to phenotypic data, to provide an unbiased estimate of genetic diversity. Although it is tempting to pick the most diverse isolates for in-depth genomic characterization, this makes it more difficult to compare results among studies (37). The level of genetic diversity we uncovered in the vaginal yeast populations suggests a relatively high level of standing genetic variation, particularly since the available comparable WGS studies in other contexts purposefully selected isolates that maximized phenotypical differences (i.e., (21, 34). Interestingly, the presence of vaginal genetic diversity is consistent with one of the earliest genetic studies, which used DNA fingerprinting to examine diversity at a single time point in up to 14 vaginal isolates from six RVVC populations (81).

Our work establishes the importance of sample size in comparing among studies. To our knowledge, our intra-population WGS characterization of up to 12 isolates at two body sites is the largest number of isolates examined to date in any context for *C. albicans* or *N. glabratus*. Previous work in other taxonomic groups suggested that a sample size of four to eight isolates is sufficient to accurately estimate population genetics parameters such as expected heterozygosity, observed heterozygosity and pairwise genetic differentiation (i.e., F_ST_) (35–37). Our bootstrap results suggest that others interested in calculating diversity should sequence six isolates per population, given the tradeoff between cost and precision; more isolates will always be better, yet we found diminishing returns. Our results also highlight that if the goal is to compare among studies, it is vital to ensure the same number of isolates are used for diversity calculations. We hope that this work will act as a launching point for studies comparing genetic diversity among different body sites and research aims.

All three *C. albicans* RVVC populations were in a subgroup in the global phylogeny in clade 1. This subgroup is enriched for vaginal isolates compared to the entire tree and even compared to the other part of clade 1. Compared to different clades, a greater proportion of clade 1 isolates have previously been noted to be significantly associated with superficial infections (82), including in the context of R/VVC (11, 30, 31, 83). Although vaginal isolates can be found throughout the phylogenetic tree, there may be something unique to the common ancestor of the clade 1 subgroup that makes them more amenable to colonizing and invading epithelial surfaces in general and hence able to cause vaginal diseases (84, 85). Most phenotypes of potential clinical interest have previously been found to vary among isolates within the same clade, precluding clear phenotype × clade associations (e.g., (86, 87)).

However, it may be that more fine-scale phylogenetic resolution is required to tease apart relationships; if only a subgroup of clade 1 isolates is enriched for a particular phenotype, that might not be seen if all clade 1 isolates are grouped together. Sala *et al.* recently found that VVC isolates induced greater fungal shedding from epithelial cells and differently stimulated epithelial signaling pathways compared to isolates from healthy women (88). This is the clearest *in vitro* assay able to differentiate VVC and healthy isolates phenotypically, and hence, a strong target for a GWAS analysis to potentially pinpoint the genetic basis of this seemingly important trait. It will be of great interest in the future to determine whether there is a genotypic association between common variants in the subgroup of clade 1 isolates (including isolates from other body sites) and their interaction with vaginal epithelial cells.

Significantly less work has been done to examine *N. glabratus* in the context of R/VVC compared to *C. albicans*, despite the increasing incidence of *N. glabratus* globally as an etiological agent of R/VVC (12, 89). We found only a single vaginal isolate out of 526 total isolates with WGS data on NCBI. This sharply contrasts with *C. albicans*, where 35 % of all sequenced isolates in the current phylogeny have been annotated as vaginal origin (20). Surprisingly, vaginal isolates form over 10% of the isolates in the *N. glabratus* MLST database (90). Those isolates are widely distributed among ST groups. This highlights a gap in inclusion of vaginal isolates in *N. glabratus* studies that use WGS.

## Conclusion

We have conducted the most extensive study to date that employed whole genome sequencing, modern methods for calculating and comparing diversity, and high throughput phenotypic analyses to compare vaginal and rectal isolates from participants with a history of symptomatic RVVC. We find no evidence that rectal isolates are different than vaginal isolates, which is inconsistent with the hypothesis that the GI tract is a source population for vaginal reinfection. We observed a near-identical average nucleotide diversity between our populations and some populations from commensal (*C. albicans*) and BSI (*N. glabratus*) settings. It remains unanswered how the nucleotide diversity is kept at such elevated levels despite repeated use of antifungals by individuals with RVVC. Combined, this study provides relevant information on the baseline diversity of RVVC vaginal and rectal populations. It emphasizes the need to investigate the role of other body sites in shaping fungal microbial communities across various contexts.

## Data availability

FASTQ files generated for this project have been deposited at the National Center for Biotechnology Information (NCBI) Sequence Read Archive under BioProject ID PRJNA991137. All phenotypic data and code required to reproduce figures, and statistical analyses are available at https://github.com/acgerstein/THRIVE_yeast-VR. Large files (e.g., BAM, VCF) are available upon request.

## Acknowledgements & funding

We are extremely thankful to the THRIVE-yeast participants. We are grateful to Dr. Alicia Berard, Dr. Adam Burgener, Kenzie Birse and other THRIVE investigators for sharing their knowledge and resources, which were integral in the initiation of this study and thank other members of the THRIVE-yeast research team including study clinicians, laboratory and administrative staff for their many contributions. The study was funded through the Manitoba Medical Service Foundation, a University of Manitoba Faculty of Science Collaborative grant, the Canadian Institute for Advanced Research (CIFAR), and the National Science and Engineering Rearch Council of Canada. ACG acknowledges the support of the CIFAR Azrieli Global Scholars Program and start-up funding from the University of Manitoba. A-RAB and AS were supported by EvoFunPath (NSERC CREATE) fellowships. DAH was funded by a University of Manitoba Research Award and the University of Manitoba Department of Microbiology Fellowship for Education Purposes. AdG was supported by a University of Manitoba Science Students’ Association and the Faculty of Science Undergraduate Student Research Award (USRA). BM was supported by a Faculty of Science USRA. Some swabs and Amies buffer tubes were donated by COPAN Italia.

