## Supplemental Figures for "Migration and standing variation in vaginal and rectal yeast populations in recurrent vulvovaginal candidiasis"

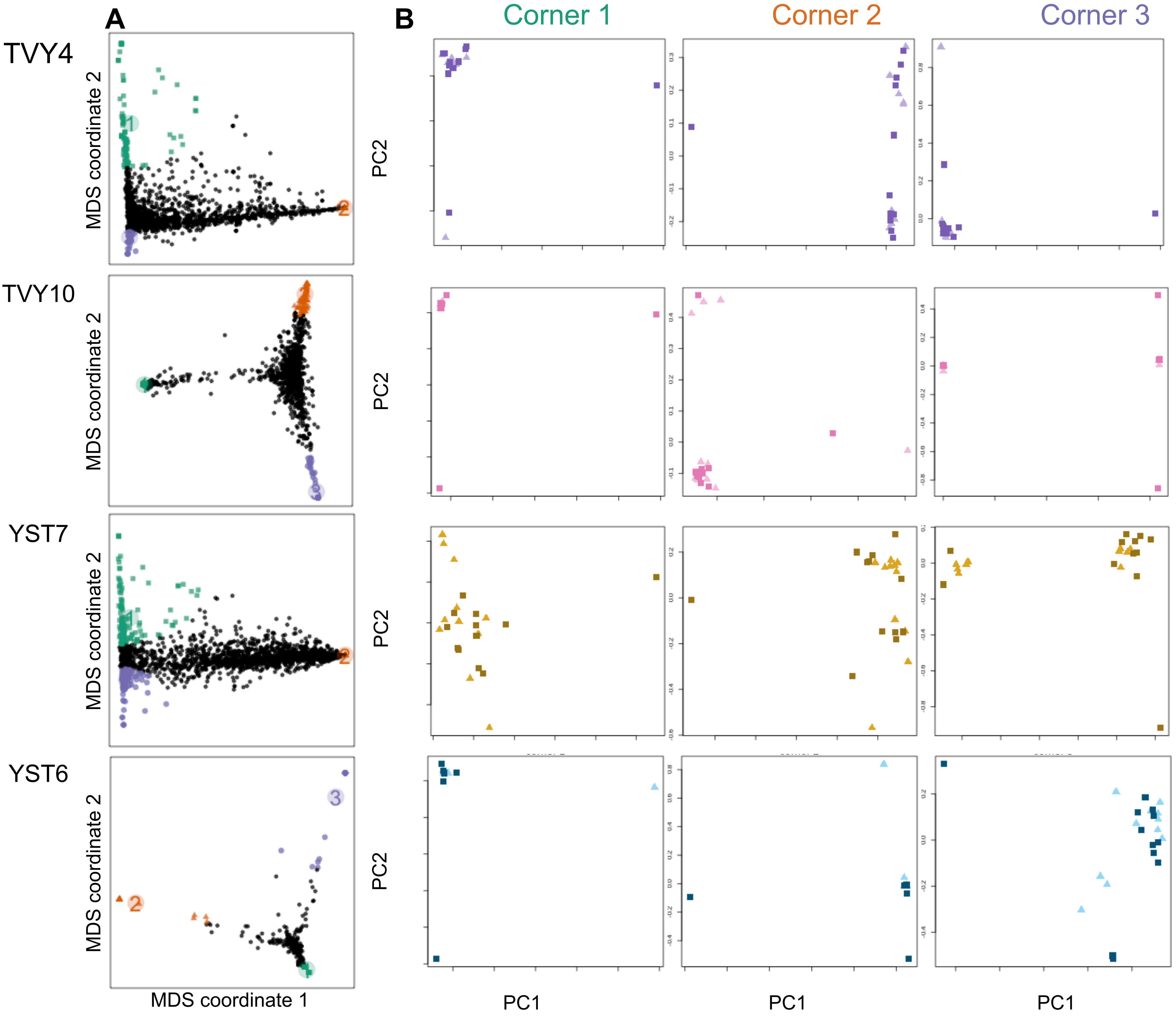


**Fig. S1** Local PCA constructed with the regions of heterogeneity in 5kb windows across the genome. Three groups of regions of high heterogeneity (A) were used in generating the PCA plot (B). This further highlights the overlapping relationship observed in the phylogeny.

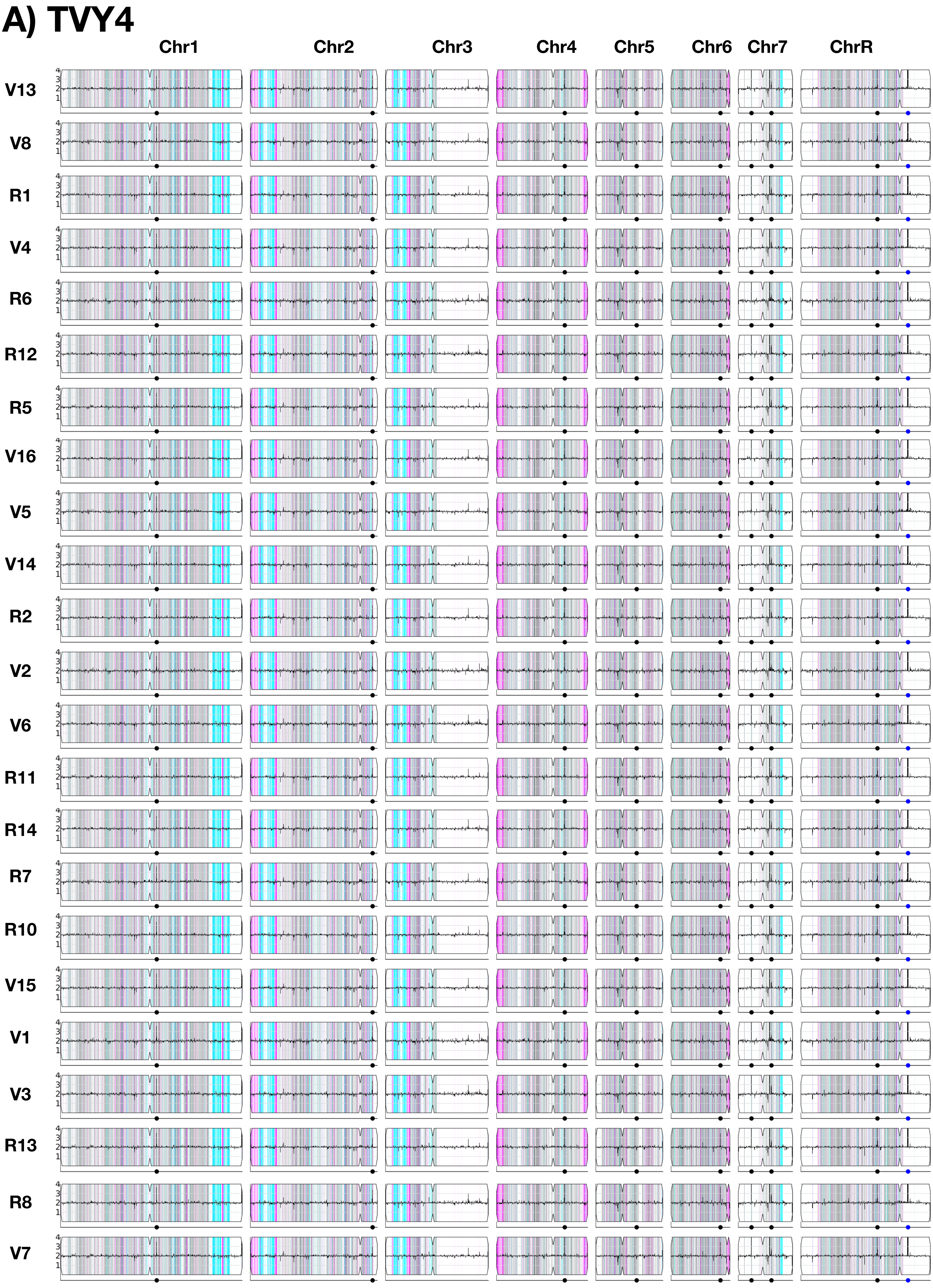
A) TVY4
Chr1
Chr2
Chr3
Chr4
Chr5
Chr6
Chr7
ChrR
V13
V8
R1
V4
R6
R12
R5
V16
V5
V14
R2
V2
V6
R11
R14
R7
R10
V15
V1
V3
R13
R8
V7

B) TVY10

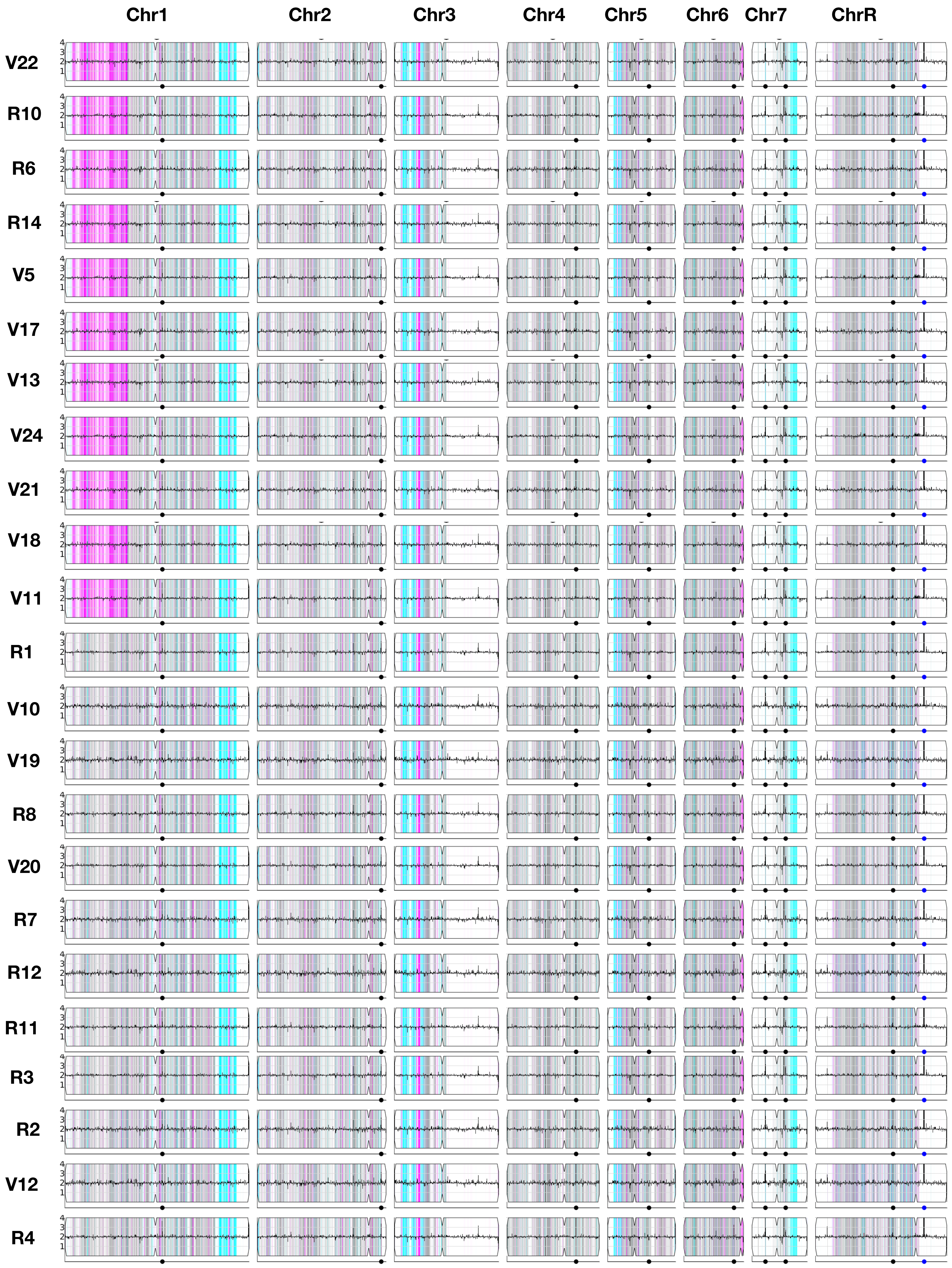

Chr1
Chr2
Chr3
Chr4
Chr5
Chr6
Chr7
ChrR
V22
R10
R6
R14
V5
V17
V13
V24
V21
V18
V11
R1
V10
V19
R8
V20
R7
R12
R11
R3
R2
V12
R4

### C) YST7

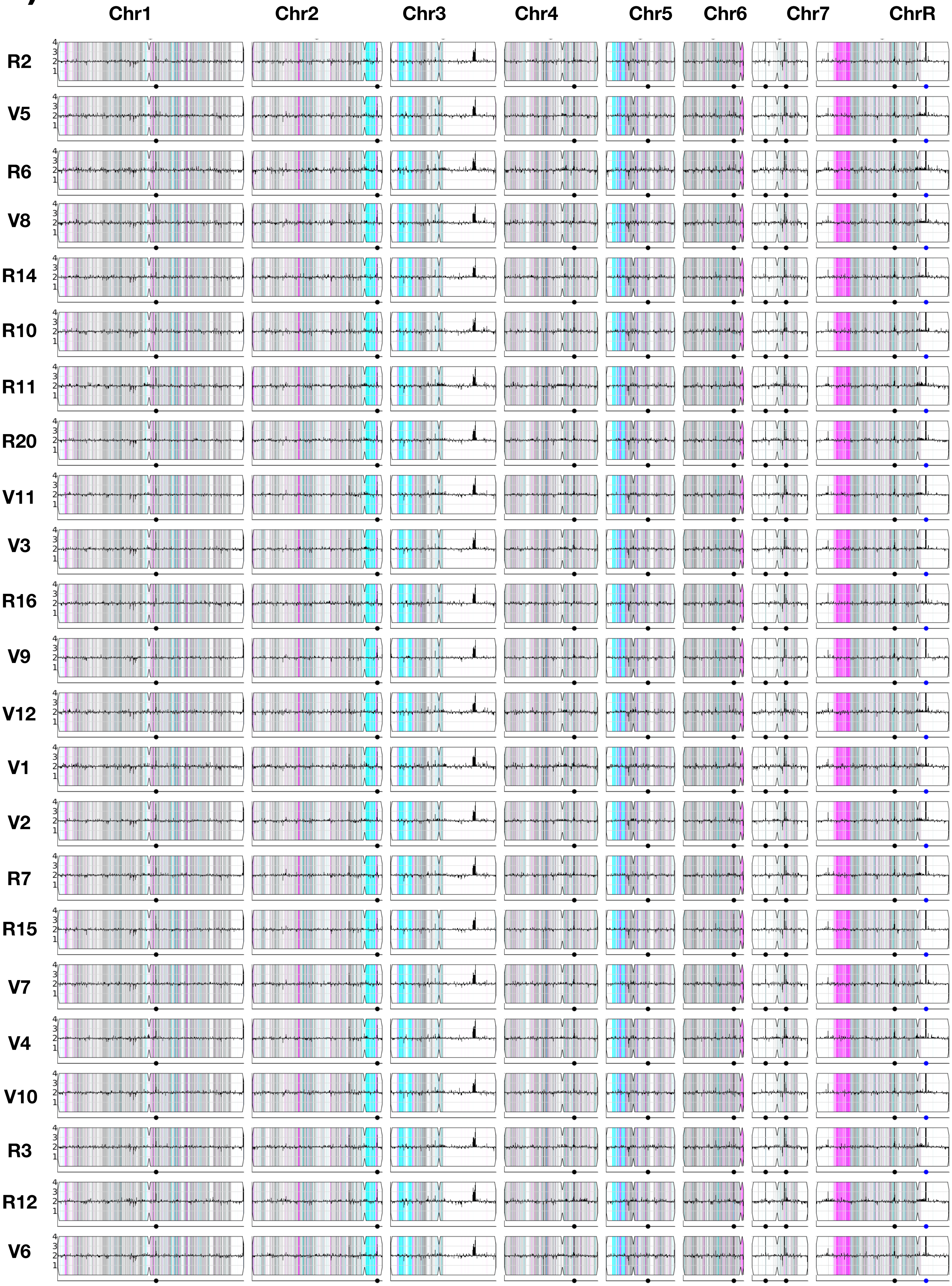
Chr1
Chr2
Chr3
Chr4
Chr5
Chr6
Chr7
ChrR
R2
V5
R6
V8
R14
R10
R11
R20
V11
V3
R16
V9
V12
V1
V2
R7
R15
V7
V4
V10
R3
R12
V6

**Fig. S2** Copy number variation and loss of heterozygosity were quantified in comparison to the SC5314 A21 reference genome using YMAP (Abbey *et al*., 2014) from isolates from (A) TVY4, (B) TVY10, and (C) YST7. The order of isolates matches that of the phylogenies shown in Figure 2. The density of single nucleotide polymorphisms is displayed as vertical lines along the length of each chromosome. Colour indicates the frequency of SNPs in 5 kb bins relative to the ancestor: white is homozygous in both the ancestror and the isolate of interest, heterozygous SNPs are shown in grey, and homozygous SNPs are color-coded to indicate the retained homolog, with cyan for “AA” and magenta for “BB”. Copy number in each bin is depicted by the black line, with the y-axis representing relative copy number; a region with increased reads is visualized as an upward spike in the region. Centromere locations are represented as indentations on the top and bottom of each chromosome box, and the dots along the bottom line mark the positions of major repeat sequences.

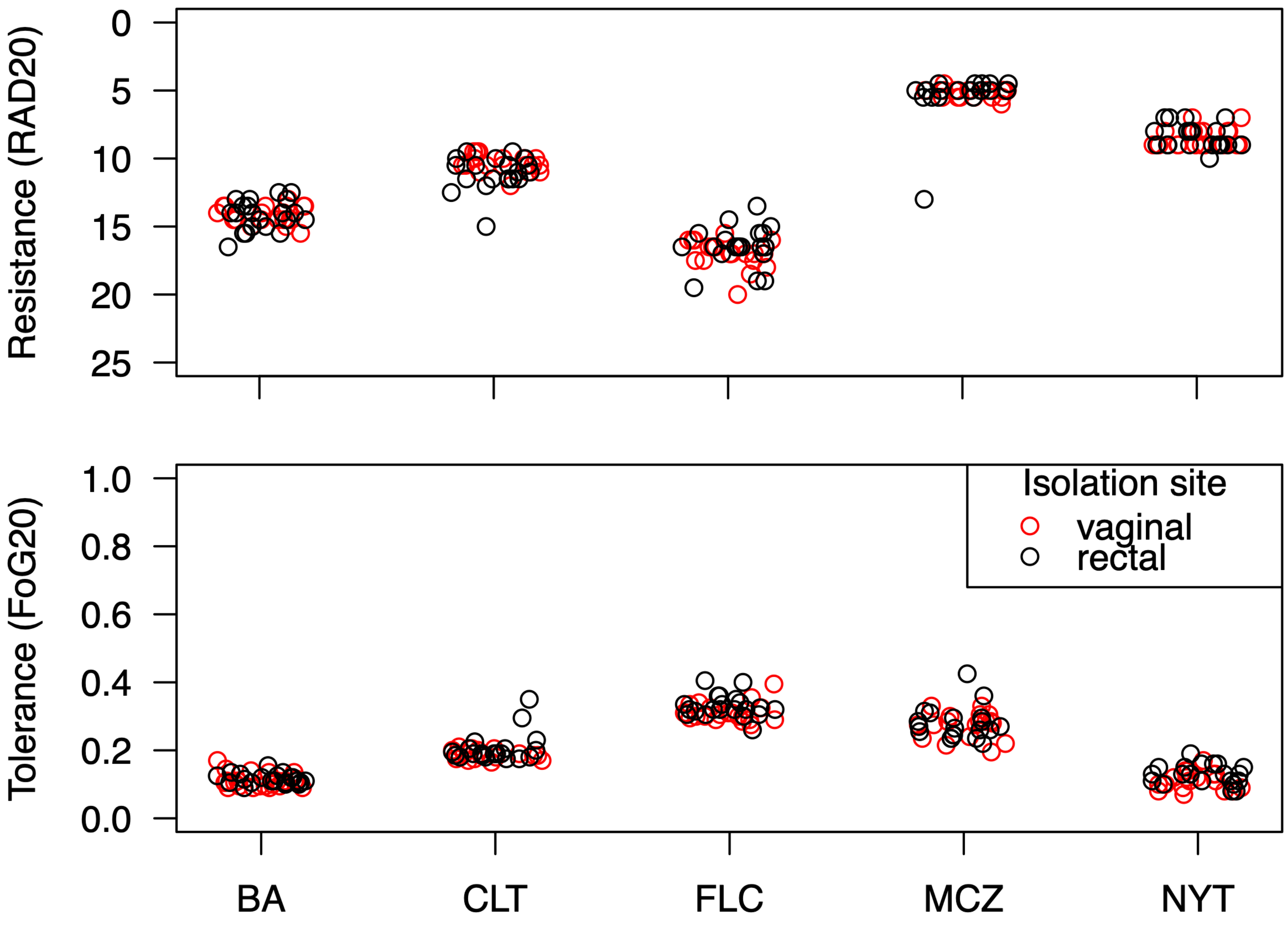


**Fig. S3** Drug response (top: resistance, bottom: tolerance) of 12 vaginal and 12 rectal isolates from YST7 C. albians isolates to five different drugs (FLC: fluconazole, CLT: clotrimazole, MCZ: miconazole, NYT: nystatin, BA: boric acid).

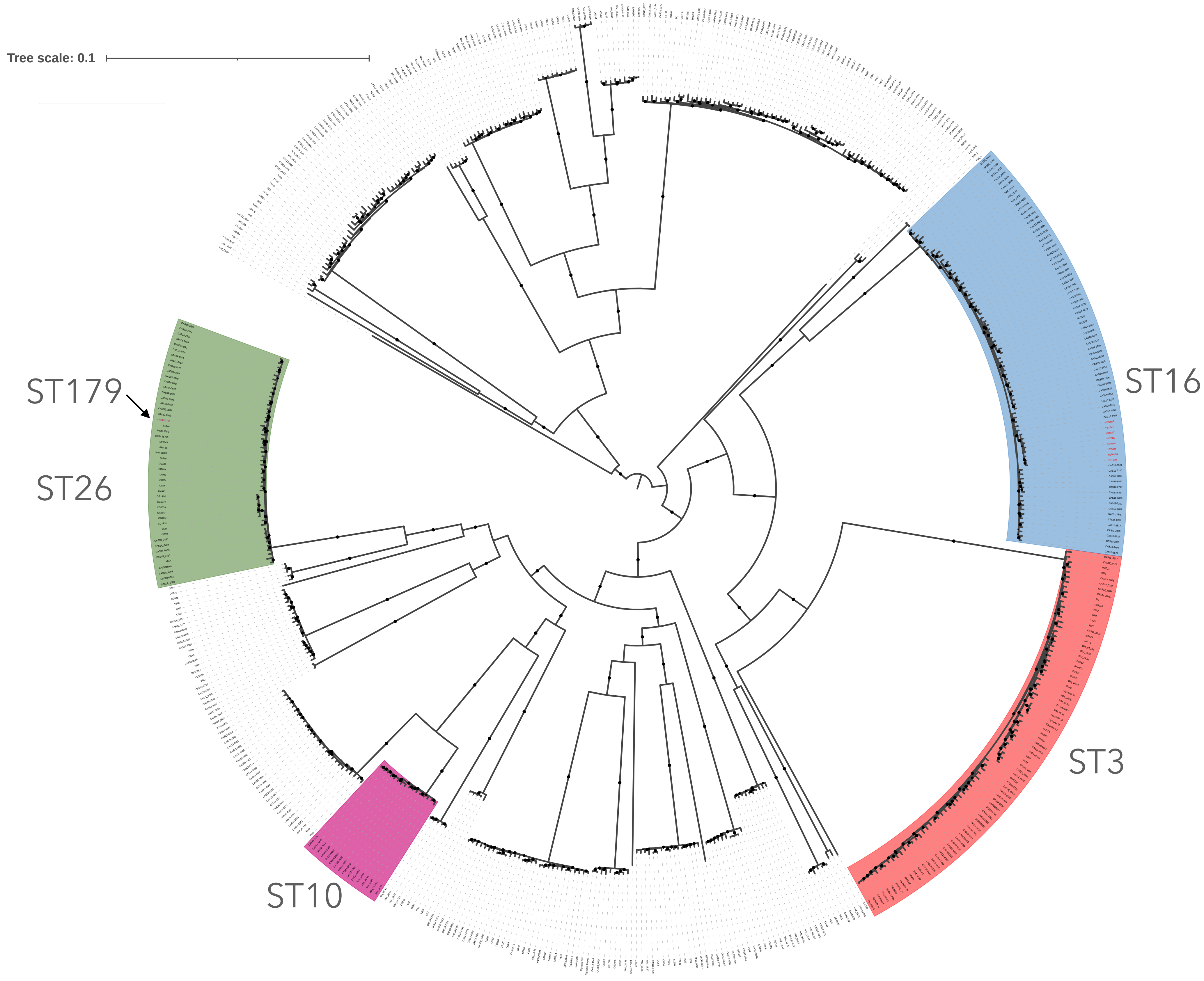


**Fig. S4** Relationship among *N. glabratus* isolates included in genetic variation comparision.
